# Removing the Gβγ-SNAP25 brake on exocytosis enhances insulin action, promotes adipocyte browning, and protects against diet-induced obesity

**DOI:** 10.1101/2020.04.29.069138

**Authors:** Ryan P. Ceddia, Zack Zurawski, Analisa Thompson Gray, Feyisayo Adegboye, Ainsley McDonald-Boyer, Fubiao Shi, Dianxin Liu, Jose Maldonado, Jiesi Feng, Yulong Li, Simon Alford, Julio E. Ayala, Owen P. McGuinness, Sheila Collins, Heidi E. Hamm

## Abstract

Negative regulation of exocytosis from secretory cells throughout the body is accomplished through inhibitory signals from G_i/o_ G protein-coupled receptors by Gβγ subunit inhibition of two common mechanisms: (i) decreased calcium entry and (ii) direct interaction of Gβγ with the Soluble *N*-ethylmaleimide-sensitive factor Attachment Protein (SNAP) Receptor (SNARE) plasma membrane fusion machinery. We have previously shown that disabling the second mechanism with a truncation of SNAP25 (SNAP25^Δ3/Δ3^) decreases the affinity of Gβγ for the SNARE complex, leaving exocytotic fusion as well as modulation of calcium entry intact but disabling GPCR inhibition of exocytosis. Here we report significant beneficial metabolic remodeling in mice carrying this mutation. Chow-fed SNAP25^Δ3/Δ3^ mice exhibit enhanced insulin sensitivity and increased beiging of white fat. In response to a high fat diet, the metabolic protection was amplified in SNAP25^Δ3/Δ3^ mice. Glucose homeostasis, whole body insulin action, and insulin-mediated glucose uptake into white adipose tissue were improved along with resistance to diet-induced obesity. This metabolic protection in SNAP25^Δ3/Δ3^ mice occurred without compromising the physiological response to fasting or cold. All metabolic phenotypes were reversed at thermoneutrality, suggesting basal autonomic activity is required. Direct electrode stimulation of sympathetic neurons exocytosis from SNAP25^Δ3/Δ3^ inguinal adipose depot resulted in enhanced and prolonged norepinephrine release. Thus, the Gβγ-SNARE interaction represents a cellular mechanism that deserves further exploration as a new avenue for combatting metabolic disease.

**GRAPHICAL ABSTRACT:** 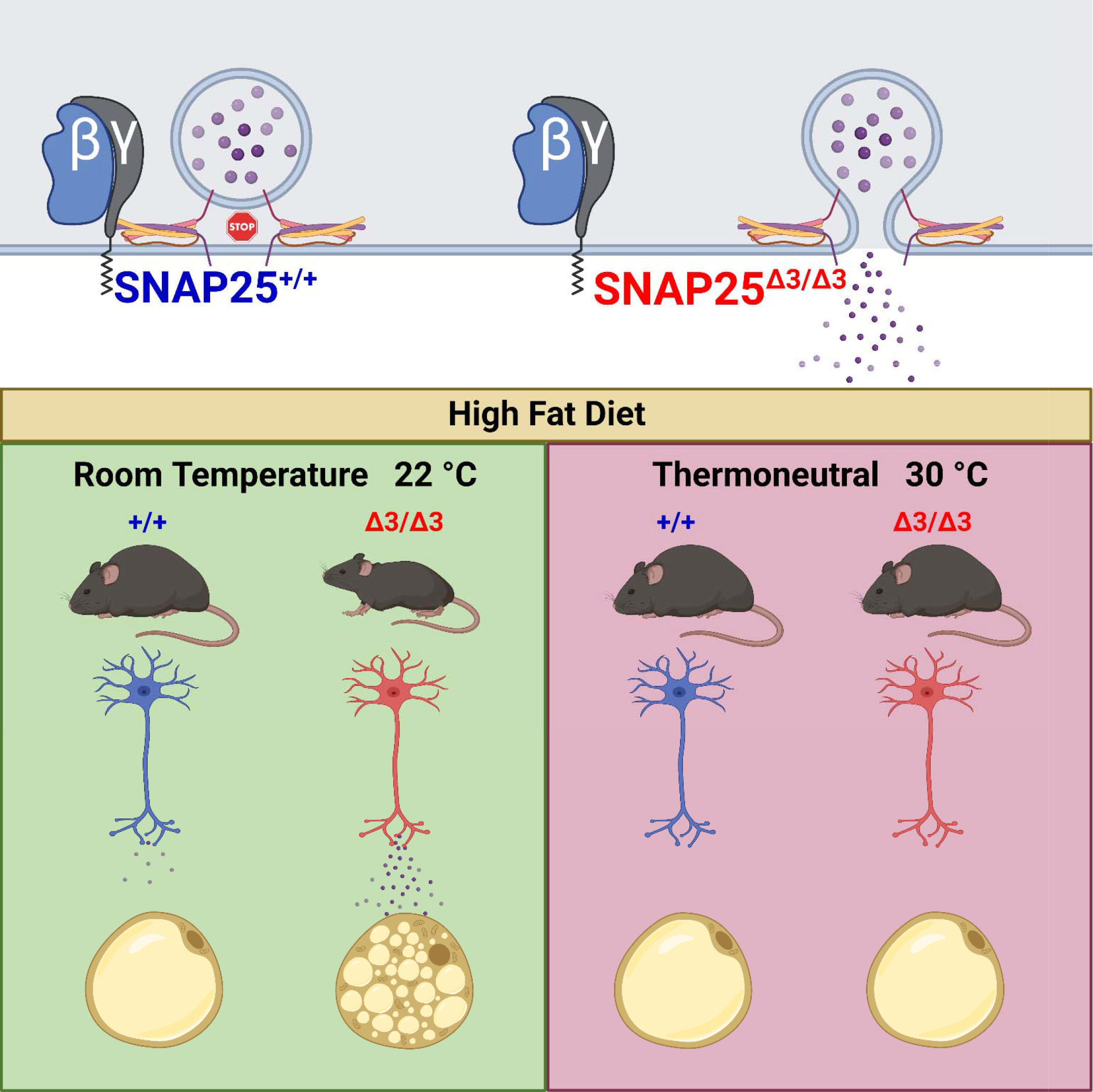

## Introduction

Metabolic diseases, such as diabetes and obesity, have an estimated annual global economic impact in the trillions of (US) dollars (1–5). Therapeutic approaches to lower metabolic risk include augmenting insulin secretion, limiting adiposity either by suppressing food intake or increasing energy expenditure, and/or by improving insulin action. G protein coupled receptors (GPCRs) in metabolically important target tissues such as the liver, brain, adipose tissue and pancreatic islets play critical roles in metabolic regulation and energy balance (6). Particularly in type 2 diabetes, pharmacologic agents have leveraged incretin-mediated GPCR signals to augment glucose-stimulated insulin secretion (GSIS) and attenuate weight gain (7). However, there may be additional approaches to manipulate GPCR signaling to treat metabolic disease.

GPCRs mediate downstream signaling events through the activation of heterotrimeric G proteins (e.g., Gα and Gβγ). Signaling events mediated by the classical Gα subunits are well known and targeting either the receptor or the Gα-generated signal in specific target tissues have been successful in many spheres of pharmacology including metabolic disease (8, 9). By contrast, signaling via the Gβγ-subunits has received less attention. They differ from Gα in their physiologic roles, which include regulation of K^+^ and Ca^+2^ ion channels via their membrane localization (10, 11). One function of Gβγ is to transduce feedback signals that modulate or inhibit neurotransmitter and hormone secretion (12). This Gβγ mechanism curbs GSIS in the pancreatic β-cell by either modulating calcium entry and/or by directly binding to the exocytotic fusion complex (13). In β-cells norepinephrine (NE) is thought to inhibit insulin exocytosis via the α_2_-adrenergic receptor (α_2_AR) through the interaction of Gβγ and SNARE proteins (13). We have shown in neurons that this inhibition requires Gβγ binding to the SNARE complex through the last three amino acids of SNAP25 (14–16). To specifically study the contribution of Gβγ to the exocytotic fusion step *in vivo*, we developed an allele of SNAP25 encoding a protein that lacks these last three amino acids (therefore called the SNAP25Δ3 protein) and created a mouse where SNAP25Δ3 replaces SNAP25 on both alleles (16). We have called this mouse SNAP25^Δ3/Δ3^. We have previously shown that SNAP25Δ3 can form SNARE complexes that undergo calcium-synaptotagmin mediated zippering and exocytosis identical to that of SNAP25, but its ability to interact with Gβγ, and the resulting GPCR-mediated inhibitory effect on exocytosis is disabled (16, 17). Using the SNAP25^Δ3/Δ3^ mouse we previously demonstrated the importance of the Gβγ-SNARE pathway in several signaling processes, including stress responses and pain processing, long-term potentiation and spatial memory (16, 18). However, the impact on metabolism and response to diet-induced obesity (DIO) are unknown.

GPCR- and SNARE-dependent signals are involved in regulating metabolically important activities, such as feeding behavior, energy balance, insulin secretion and thermoregulation (19–21). However, the specific role that Gβγ-dependent signals play in regulating these physiologic responses are unknown. For example, when a palatable high fat diet (HFD) is made available, it is unknown if Gβγ-SNARE-dependent signals serve to limit or amplify weight gain or contribute to glucose dysregulation and metabolic inflexibility commonly seen in metabolic disease (22, 23). As Gβγ-SNARE- dependent signals can serve as physiological brakes, they may restrain insulin secretion, impair insulin action, limit satiety or attenuate energy expenditure, thus aggravating adiposity and impairing glucose homeostasis. We hypothesized that attenuation of this physiological brake using the SNAP25^Δ3/Δ3^ mouse would improve glucose homeostasis and improve metabolic disease phenotypes in response to a palatable HFD. Our data described here demonstrate that chow-fed SNAP25^Δ3/Δ3^ mice are more insulin sensitive and display increased adipose tissue beiging. Moreover, in response to a HFD challenge, the removal of this Gβγ-SNARE braking mechanism attenuates adiposity, improves insulin action, and amplifies sympathetic activity and subsequent browning of adipose tissue. Reducing sympathetic tone by placing animals in a thermoneutral environment removes the metabolic protection afforded by this Gβγ-SNARE braking mechanism. The resistance to DIO and improved glucose homeostasis when this braking mechanism is disabled suggest the Gβγ-SNARE interaction is a physiologically important mechanism that may be leveraged to treat metabolic disease.

## Results

### Normal growth curves with increased white adipose tissue beiging in chow-fed SNAP25^Δ3/Δ3^ mice

Body weights of male and female SNAP25^Δ3/Δ3^ mice were monitored over a 60-week period while being fed a standard chow diet; no differences were observed in growth curves compared to their SNAP25^+/+^ littermates (**Fig. 1A; Fig. S1A**). Weights of internal organs and tissues were examined in a second cohort of 15-week-old mice. No statistically significant difference in most organ weights was observed. As there was a trend toward slightly smaller adipose depots in SNAP25^Δ3/Δ3^ mice (**Fig. 1B**), we examined adipose histology. We observed a modest increase in UCP1-positive beige adipocytes in the inguinal white adipose tissue (iWAT) depot without an accompanying change in the interscapular brown adipose tissue (**Fig. 1C,D**). Consistent with a reduction in adipose tissue mass, we observed a decrease in the amount of circulating leptin (3.01 ± 0.46 vs. 1.24 ± 0.27 ng/ml leptin; *N* = 9 SNAP25^+/+^ vs. 12 SNAP25^Δ3/Δ3^ male mice; mean ± SEM; P = 0.002). These differences in adipose weight and remodeling were more evident in female mice (**Fig. S1**). A cohort of 15-week-old SNAP25^Δ3/Δ3^ female mice weighed slightly less than SNAP25^+/+^ mice (**Fig. S1B**), had smaller fat pads (**Fig. S1C**) and adipocyte size was reduced in both iBAT and iWAT (**Fig. S1D,E**). Though lacking a strong effect on the overall size and appearance of the mice, loss of the Gβγ-SNARE interaction led to a reduction in the size of adipocytes and their fat pad depots and is associated with increased WAT beiging.

**Figure 1.**
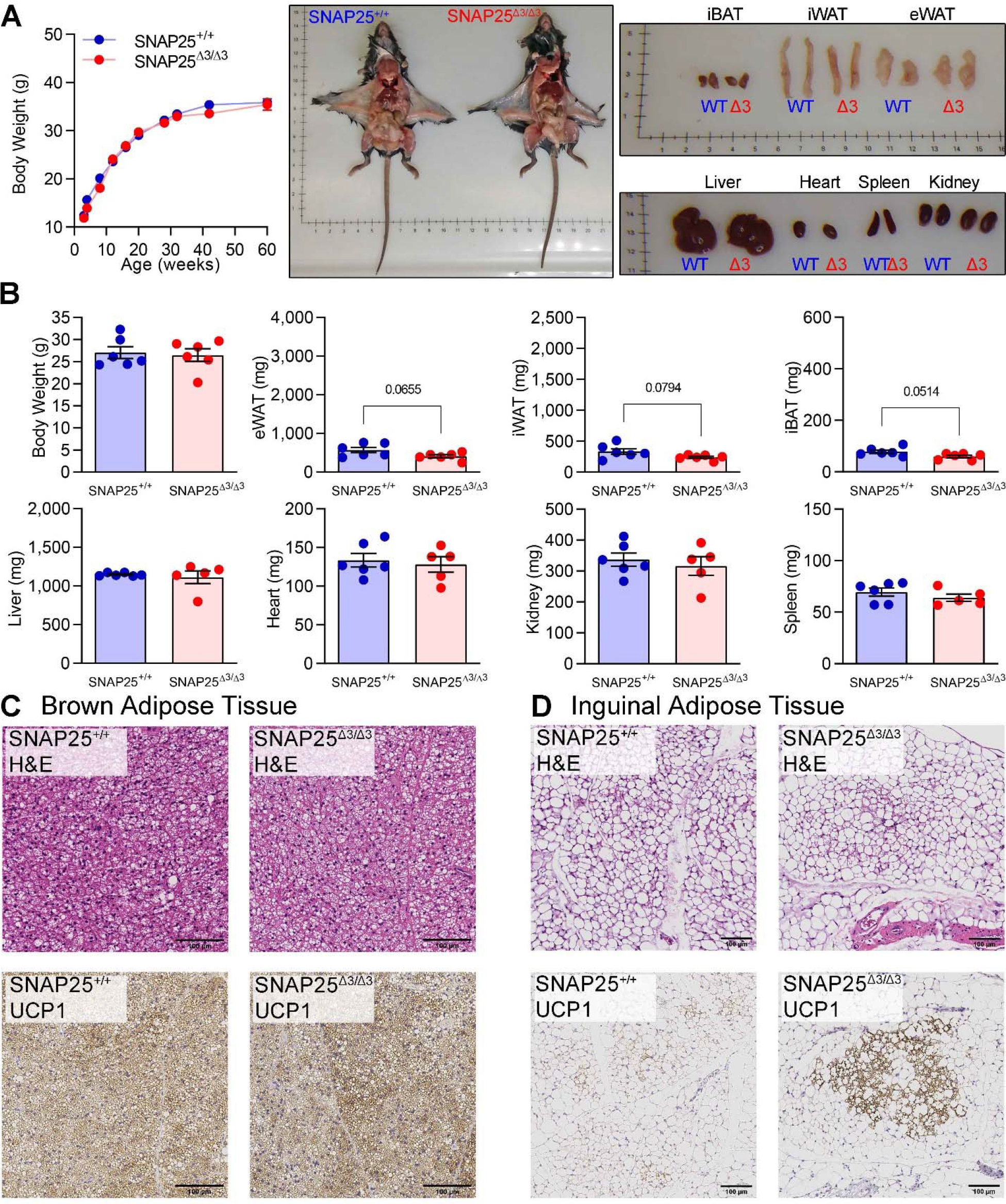
Body and tissue weights are unchanged while white adipose tissue beiging is increased in chow-fed, male SNAP25^Δ3/Δ3^ mice **A.** Body weights of SNAP25^+/+^ and SNAP25^Δ3/Δ3^ male mice were fed standard chow diets (P = 0.3849, effect of genotype; P = 0.0820, effect of genotype × time interaction). *N* = 11 SNAP25^+/+^, 11 SNAP25^Δ3/Δ3^. Analysis was performed by two-way ANOVA with repeated measures and post hoc analyses were performed using Bonferroni multiple comparisons test for SNAP25 genotype only. **B.** Body and tissue weights of a separate cohort of chow-fed mice that were euthanized at 15-weeks of age (pictured). *N* = 6 SNAP25^+/+^, 6 SNAP25^Δ3/Δ3^. Analyses were performed using t-test. **C.** Representative H&E- and UCP1-stained sections of iBAT. **D.** Representative H&E- and UCP1-stained sections of iWAT. Images are a representative sample from three mice from each group. Values are expressed as mean ± SEM.

### Lower glucose stimulated insulin secretion in vivo without defects in glucose regulation suggests improved insulin action in SNAP25^Δ3/Δ3^ mice fed chow diet

To determine if disabling the Gβγ inhibition of SNARE-mediated exocytosis would improve glucose homeostasis in chow-fed mice, we measured circulating glucose and insulin during an intraperitoneal glucose tolerance test in 14-week-old chow-fed mice. Glucose tolerance in the SNAP25^Δ3/Δ3^ mice was not different from SNAP25^+/+^ mice (**Fig. 2A**). While basal insulin levels in SNAP25^Δ3/Δ3^ mice were unaltered, insulin concentrations were reduced during the GTT compared to SNAP25^+/+^ mice (**Fig. 2B**) consistent with an improved insulin action. Since the incretin effect is bypassed with IP injection of glucose, we performed an oral glucose tolerance test in a separate cohort of chow-fed mice to test whether this approach might unmask an augmentation of insulin secretion in the SNAP25^Δ3/Δ3^ mice (24). Oral glucose tolerance was again not different between genotypes (**Fig. 2C**). However, as was observed with the IP route of delivery, plasma insulin levels were lower in SNAP25^Δ3/Δ3^ mice after the glucose gavage (**Fig. 2D**). The lower levels of insulin with a normal glucose tolerance are consistent with improved insulin sensitivity in SNAP25^Δ3/Δ3^ mice.

**Figure 2.**
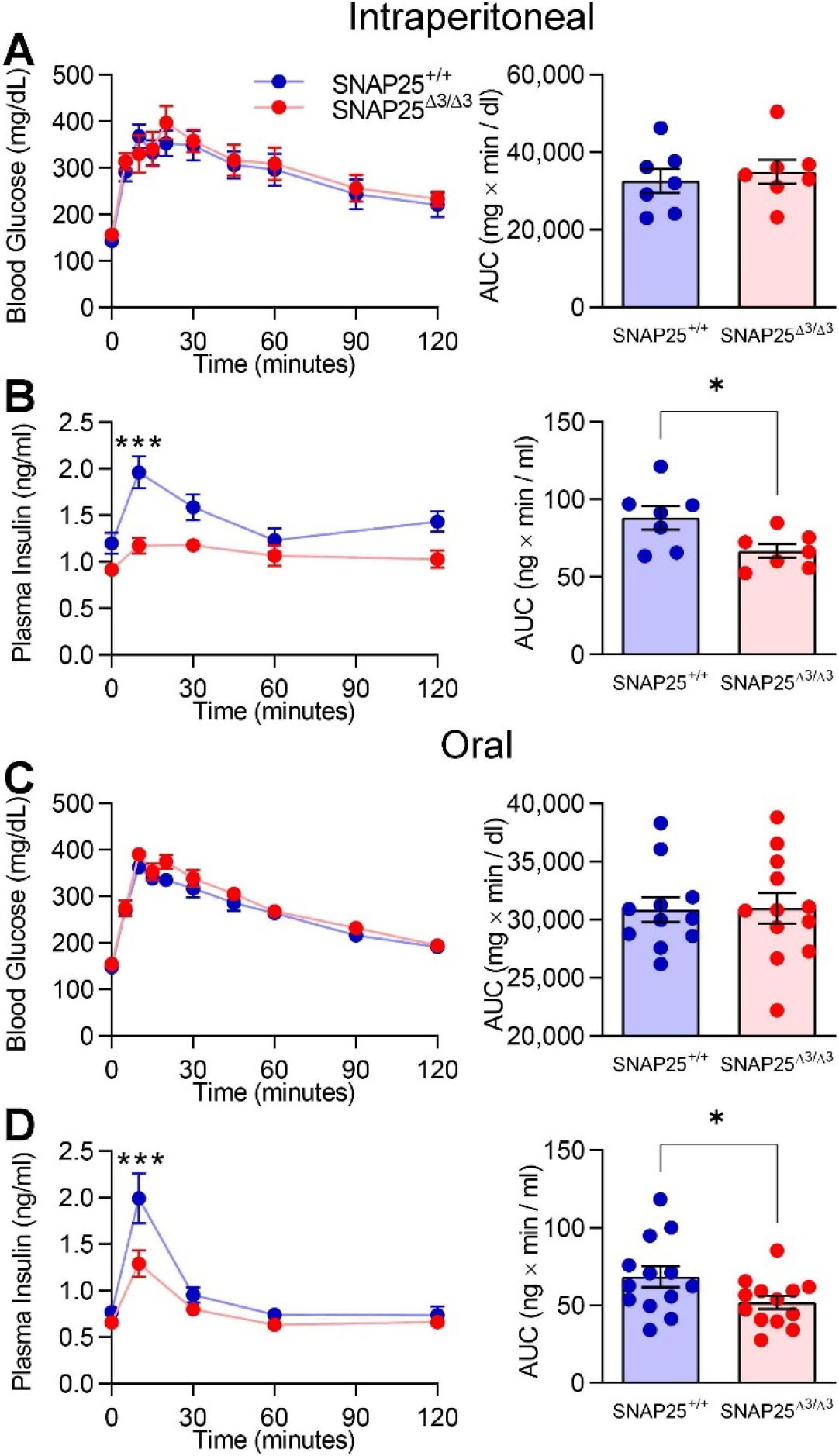
**Glucose tolerance is normal but circulating insulin is reduced in chow-fed SNAP25**^Δ**3/**Δ**3**^ **mice A.** Blood glucose and **B.** plasma insulin levels (P = 0.0176, effect of genotype; P = 0.0031, effect of genotype × time interaction) during an IP-GTT in chow-fed male SNAP25^+/+^ and SNAP25^Δ3/Δ3^ at 14 weeks of age. *N* = 8 SNAP25^+/+^, 8 SNAP25^Δ3/Δ3^. **C.** Blood glucose and **D.** plasma insulin levels (P = 0.0730, effect of genotype; P = 0.0009, effect of genotype × time interaction) during an oral-GTT in chow-fed male SNAP25^+/+^ and SNAP25^Δ3/Δ3^ at 15 weeks of age. *N* = 13 SNAP25^+/+^, 13 SNAP25^Δ3/Δ3^. * p<0.05. Values are expressed as mean ± SEM. For glucose and insulin time course, two-way ANOVAs were performed with repeated measures and post hoc analyses were performed using Bonferroni multiple comparisons test for SNAP25 genotype only and are indicated on figures. Areas Under the Curve (AUC) were analyzed by t-test.

There was no augmentation of insulin secretion during the GTT *in vivo* (**Fig. 2B,D**), although this could be explained by an accompanying improvement in insulin action. Other investigators reported in an immortalized mouse β-cell line that inhibition of insulin exocytosis by NE via α_2_AR signaling was mediated by the Gβγ-SNARE interaction (13). We sought to determine in freshly isolated perifused islets if loss of the Gβγ-SNARE interaction improved GSIS and/or impaired α_2_AR dependent inhibition of GSIS. We used a selective α_2_AR agonist brimonidine (Br) to test this interaction. As shown in **Fig. 3A** and **3B**, GSIS in isolated islets from SNAP25^Δ3/Δ3^ mice was not greater than from SNAP25^+/+^ mice. While islets from SNAP25^Δ3/Δ3^ mice seemed to secrete less insulin, when multiple experiments are compiled there is no significant difference (**Fig. 3C**). Moreover, contrary to the earlier studies in the cell line (13), Br was able to equally suppress GSIS in islets from SNAP25^Δ3/Δ3^ and SNAP25^+/+^ mice (**Fig. 3A, C**). The insulin content of the islets was also not significantly affected by SNAP25 genotype (**Fig. 3D**). This indicates that α_2_AR-mediated signaling in *ex vivo* mouse islets is able to inhibit GSIS via a mechanism that is independent of the Gβγ-SNARE interaction.

**Figure 3.**
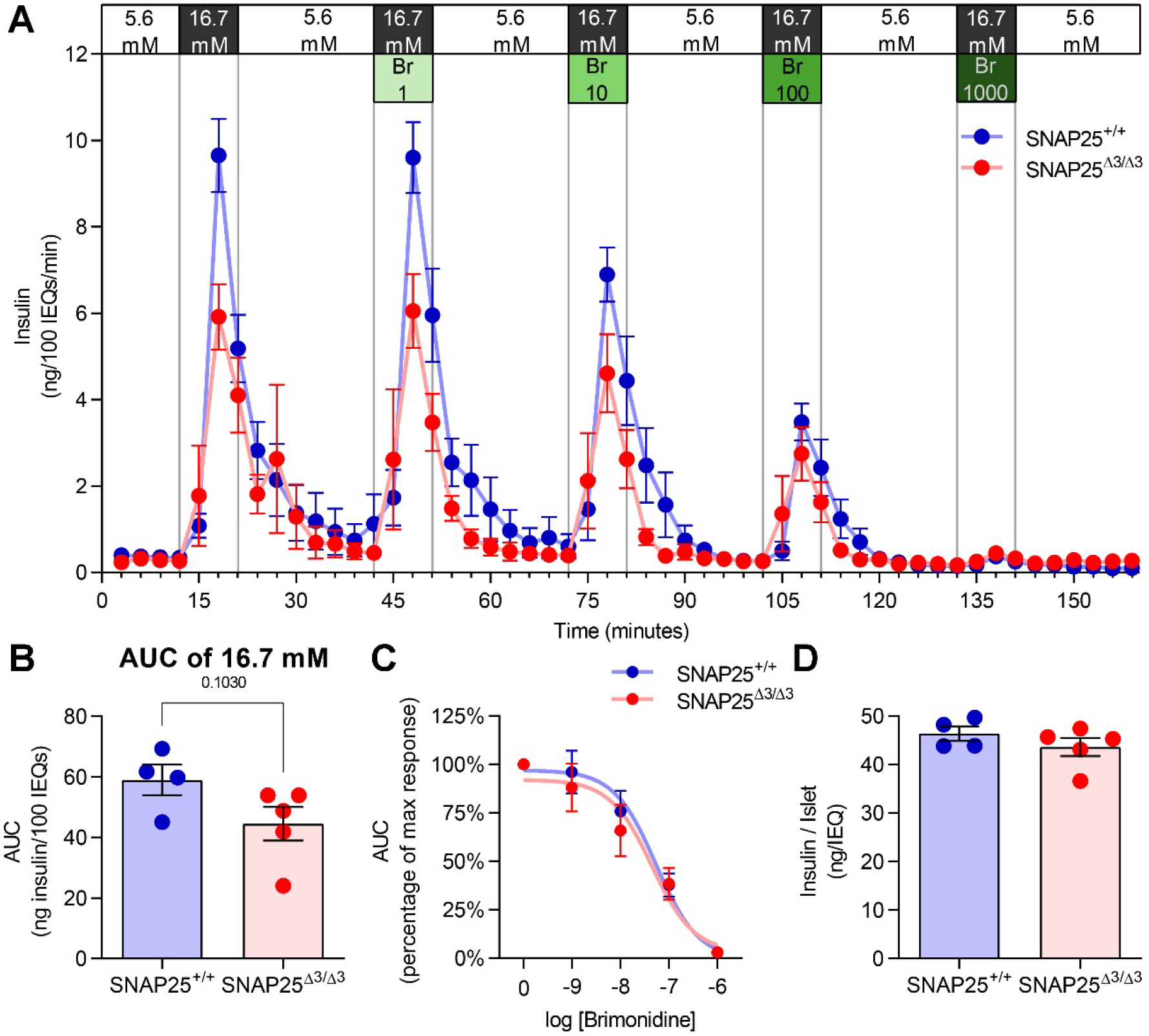
**Glucose stimulated insulin secretion is not increased in islets from SNAP25**^Δ**3/**Δ3^ mice *ex vivo* and inhibition by α**_2_AR agonist is not affected by disruption of the G**βγ**-SNAP25 interaction A.** Perifusion of islets from 12-week-old male SNAP25^+/+^ and SNAP25^Δ3/Δ3^ mice. **B.** Comparison of glucose stimulated insulin secretion (GSIS) between islets from SNAP25^+/+^ and SNAP25^Δ3/Δ3^ mice. **C.** AUC values were normalized to the individual’s maximal GSIS and a dose-response curve for the inhibition of GSIS by the α_2_AR selective agonist, brimonidine (Br), was generated. The log IC50’s were similar, being -7.262 for SNAP25^+/+^ and -7.348 for SNAP25^Δ3/Δ3^. **D.** Islet insulin content in islets from SNAP25^+/+^ and SNAP25^Δ3/Δ3^ mice. For all figures *N* = 4 SNAP25^+/+^, 5 SNAP25^Δ3/Δ3^. Values are expressed as mean ± SEM. Analyses were performed using t-test.

### Energy expenditure at room temperature and in response to cold did not differ between SNAP25 genotypes on a chow diet

We assessed if disabling the Gβγ-SNARE interaction altered the response to physiologic stressors, since we had previously reported that SNAP25^Δ3/Δ3^ mice had an altered behavioral response to acute physiological stressors, specifically elevated stress-induced hyperthermia (16, 25). We performed calorimetry studies in which energy expenditure and feeding behavior in chow-fed SNAP25^+/+^ and SNAP25^Δ3/Δ3^ mice were measured at standard housing temperature (22°C). The mice were then exposed to an acute cold challenge (6°C) to assess whether they could mount a physiologic response by increasing energy expenditure and food intake. At normal housing temperature SNAP25^Δ3/Δ3^ mice had similar rates of energy expenditure (**Fig. S2A**) and food intake (**Fig. S2B**), with normal diurnal patterns for both variables. In response to a decrease in environmental temperature both SNAP25^Δ3/Δ3^ mice and their SNAP25^+/+^ littermates mounted a robust increase in energy expenditure. This was also accompanied by a greater increase in the duration of feeding activity in SNAP25^Δ3/Δ3^ mice (**Fig. S2E**). Interestingly, any changes in autonomic activity that might occur are not global. For example, in a separate cohort, cardiovascular parameters were measured at standard housing temperature using echocardiography, and no differences were observed in the SNAP25^Δ3/Δ3^ mice (**Fig. S3**). Thus, on a chow diet SNAP25^Δ3/Δ3^ mice display subtle changes in feeding behavior that overall do not impact energy balance, and the physiologic autonomic dependent response to cold is also intact.

### Protection from high fat diet-induced obesity and adiposity in SNAP25^Δ3/Δ3^ mice

Given the apparent greater insulin sensitivity of the SNAP25^Δ3/Δ3^ mice when on chow diet (**Fig. 2**) we tested whether this phenotype would persist when challenged with a HFD. We focused on male mice given the robust improvement in insulin action observed on a chow diet. Eight-week-old SNAP25^+/+^ and SNAP25^Δ3/Δ3^ mice were provided a HFD for 8 weeks. As shown in **Fig 4A**, SNAP25^+/+^ mice rapidly gained weight on the HFD, whereas SNAP25^Δ3/Δ3^ mice were markedly resistant to HFD- induced weight gain. Body composition analyses revealed that the reduced weight gain in SNAP25^Δ3/Δ3^ mice was due to reduced adiposity (**Fig. 4B**) and was confirmed in postmortem analyses of these mice (**Fig 4C**). This was reflected in a decrease in plasma leptin (14.3 ± 1.32 vs. 3.53 ± 1.23 ng/ml leptin; *N* = 9 SNAP25^+/+^ vs. 7 SNAP25^Δ3/Δ3^; mean ± SEM; P < 0.001). In addition to a reduction in the weight of all fat depots (epididymal WAT; eWAT, inguinal WAT; iWAT, retroperitoneal WAT; rpWAT, and interscapular BAT; iBAT), liver weight and hepatic steatosis were also reduced in SNAP25^Δ3/Δ3^ compared to SNAP25^+/+^mice (**Fig. 4D, E**). The size of the kidney, spleen, heart, and most skeletal muscles were unaffected (**Fig. 4E**).

**Figure 4.**
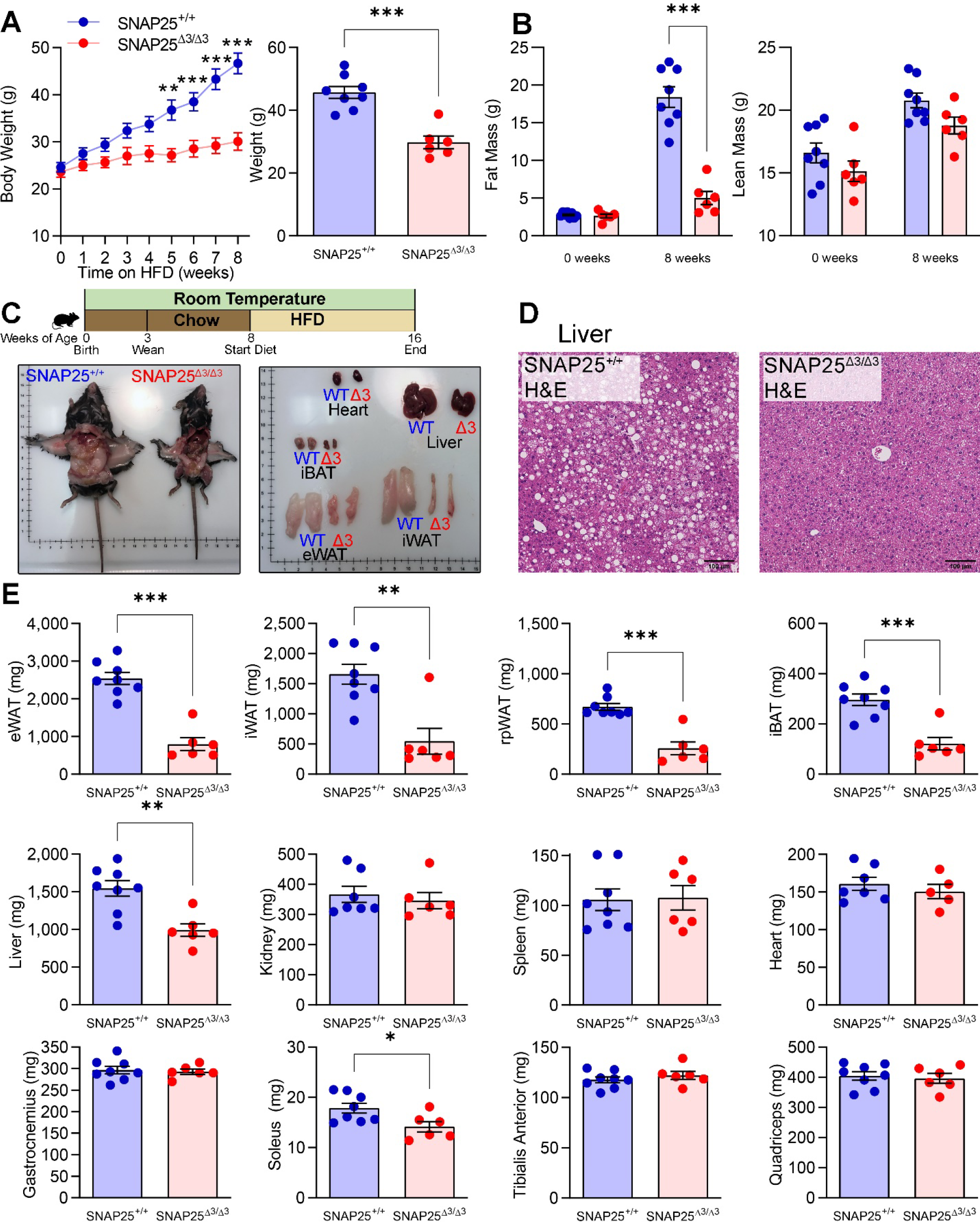
**Weight gain and adiposity induced by HFD are reduced in SNAP25**^Δ**3/**Δ**3**^ **mice A.** Male SNAP25^+/+^ and SNAP25^Δ3/Δ3^ were fed HFD for 8 weeks beginning at 8 weeks of age (P = 0.0030, effect of genotype; P < 0.001, effect of genotype × time interaction). Comparison of terminal body weights **B.** Body composition of fat mass (P < 0.001, effect of genotype; P < 0.001, effect of genotype × time interaction) and lean mass (P = 0.0891, effect of genotype) at beginning and end of HFD feeding period. **C.** Representative images of HFD-fed SNAP25^+/+^ and SNAP25^Δ3/Δ3^ mice and tissues post-mortem. **D.** Representative H&E-stained sections of livers. Images are a representative sample from three mice from each group. **E.** Tissue weights of SNAP25^+/+^ and SNAP25^Δ3/Δ3^ mice at the end of the HFD feeding period. For all figures *N* = 8 SNAP25^+/+^, 6 SNAP25^Δ3/Δ3^. * p<0.05, ** p<0.01, *** p<0.001. Values are expressed as mean ± SEM. For A (left) and B, two-way ANOVAs were performed with repeated measures and post hoc analyses were performed using Bonferroni multiple comparisons test for SNAP25 genotype only and are indicated on figures. Analyses for terminal body weight (A right) and D were performed using t-test.

### Improvement in insulin action in SNAP25^Δ3/Δ3^ mice following high fat diet challenge

Having observed a profound difference in weight gain and adiposity in SNAP25^Δ3/Δ3^ mice when challenged with a HFD, we hypothesized that glucose homeostasis would also be improved. Indeed, HFD-fed SNAP25^Δ3/Δ3^ mice showed a significant improvement in the IP-GTT compared with SNAP25^+/+^ mice (**Fig. 5A**). SNAP25^Δ3/Δ3^ mice had lower 5 h fasting glucose levels (**Fig. 5B**), congruent with their lower IP-GTT curves. Plasma insulin was also significantly lower (**Fig. 5B**). Together, these data indicate that there is a *bona fide* improvement in insulin sensitivity of the SNAP25^Δ3/Δ3^ mice.

**Figure 5.**
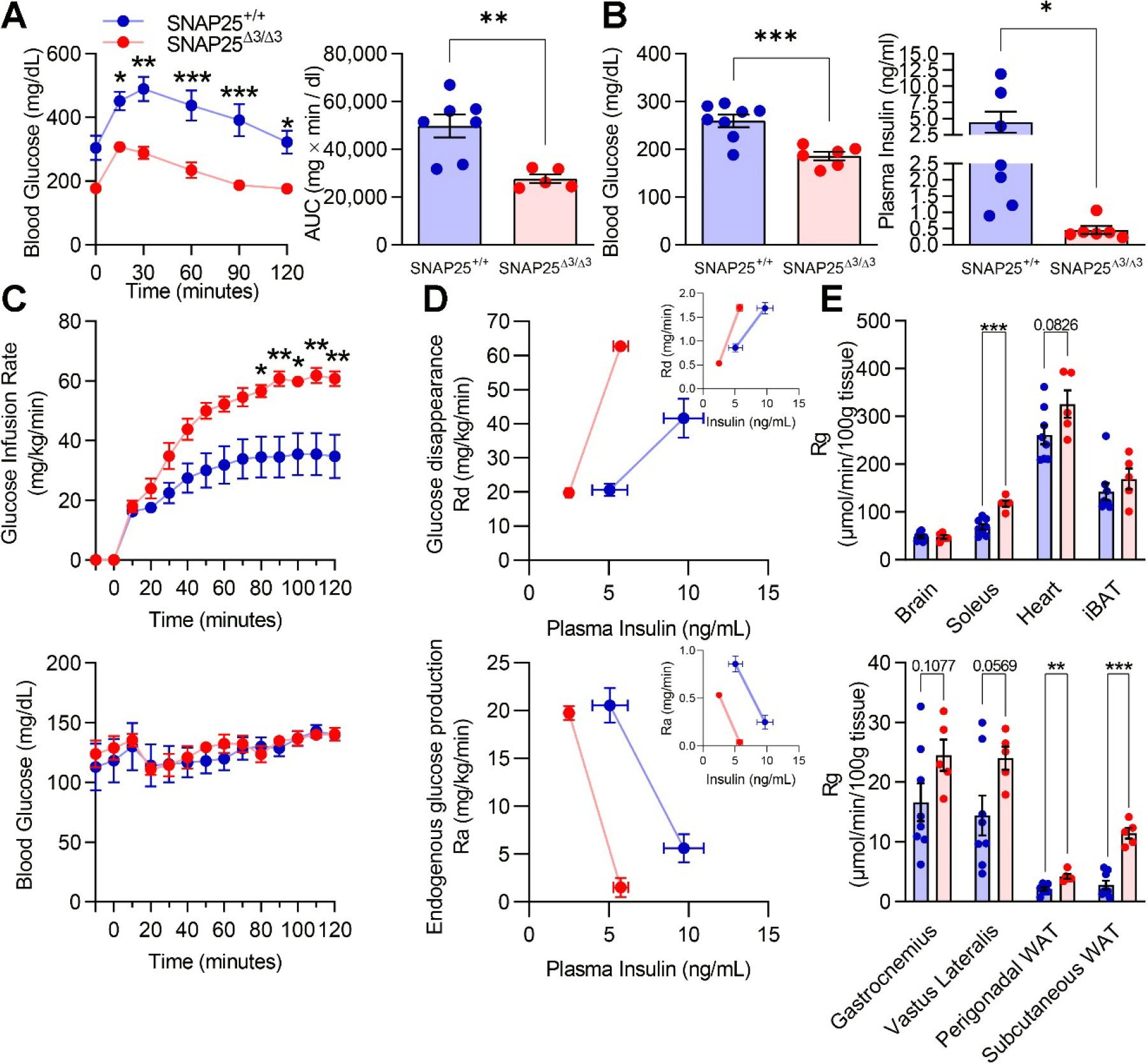
**Insulin action and glucose homeostasis during HFD challenge are improved in SNAP25**^Δ**3/**Δ**3**^ **mice A.** IP-GTT in male SNAP25^+/+^ and SNAP25^Δ3/Δ3^ mice fed HFD for 8 weeks (P = 0.0040, effect of genotype; P = 0.0424, effect of genotype × time interaction). *N* = 7 SNAP25^+/+^, 5 SNAP25^Δ3/Δ3^. **B.** Fasting blood glucose and plasma insulin levels at euthanasia. *N* = 7-8 SNAP25^+/+^, 6 SNAP25^Δ3/Δ3^. **C.** Glucose infusion rate (P = 0.0224, effect of genotype; P < 0.001, effect of genotype × time interaction) and blood glucose levels of HFD-fed SNAP25^+/+^ and SNAP25^Δ3/Δ3^ male mice during hyperinsulinemic euglycemic clamp. **D.** Glucose disappearance (Rd; mg·kg^-1^·min^-1^) and endogenous glucose production (Ra; mg·kg^-1^·min^-1^) plotted against plasma insulin levels. In inset is Ra and Rd (mg/min) not normalized to body weight plotted against plasma insulin levels . **E.** Tissue [^14^C]2-deoxy-D-glucose uptake (Rg). For C-E, *N* = 8 SNAP25^+/+^, 5 SNAP25^Δ3/Δ3^. ** p<0.01, *** p<0.001. Values are expressed as mean ± SEM. For A D, and C (glucose and GIR), two-way ANOVA was performed with repeated measures and post hoc analyses were performed using Bonferroni multiple comparisons test for SNAP25 genotype only and are indicated on figures. For A (AUC) B and E, analyses were analyzed by t-test.

To determine the contribution of specific tissues to the improved glucose homeostasis of SNAP25^Δ3/Δ3^ mice, we performed hyperinsulinemic euglycemic clamps in chronically catheterized conscious mice that had been consuming the HFD for 8 weeks (40.9 ± 2.5 vs. 27.2 ± 0.8 g body weight; SNAP25^+/+^ vs. SNAP25^Δ3/Δ3^; mean ± SEM). We determined if insulin suppression of endogenous (i.e. hepatic) glucose production and/or stimulation of peripheral glucose uptake were altered. Arterial insulin levels were lower in the basal period in SNAP25^Δ3/Δ3^ (5.8 ± 1.1 vs. 2.5 ± 0.2 ng/ml; SNAP25^+/+^ vs. SNAP25^Δ3/Δ3^; mean ± SEM) and increased with insulin infusion during the clamp period (9.7 ± 1.7 vs. 5.1 ± 0.8 ng/ml; SNAP25^+/+^ vs. SNAP25^Δ3/Δ3^; mean ± SEM). Arterial blood glucose was maintained at euglycemia in both groups throughout the clamp procedure (**Fig. 5C**). The glucose infusion rate required to maintain euglycemia was increased in SNAP25^Δ3/Δ3^ despite lower clamp insulin concentrations (**Fig. 5C**). Using [3-^3^H]-glucose to assess whole body glucose flux (mg·kg^-1^·min^-1^) we determined that the increase in glucose requirements in SNAP25^Δ3/Δ3^ resulted from an increase in whole body glucose uptake (**Fig. 5D**). Basal endogenous glucose production as well as insulin suppression of endogenous glucose production were comparable, despite the fact that the rise in insulin was substantially lower in SNAP25^Δ3/Δ3^ mice (**Fig. 5D**). As SNAP25^Δ3/Δ3^ mice were substantially leaner we also calculated glucose flux (mg/min) without dividing by body weight (**Fig 5D inset**). Whole total glucose flux was not augmented during the clamp period (**Fig 5D inset**). However, there was a shift to the left in the relationship between insulin suppression of glucose production as well as stimulation of whole-body glucose uptake. Thus, irrespective of how the data are presented both hepatic and peripheral tissues of SNAP25^Δ3/Δ3^ mice were more sensitive to insulin. To determine which tissues contributed to the increase in insulin action we assessed glucose uptake (Rg) in multiple tissues during the clamp using [^14^C]2-deoxy-D-glucose (2-DG). Rg was markedly increased in SNAP25^Δ3/Δ3^ mice in multiple skeletal muscles (soleus, and vastus lateralis) and white adipose tissue depots (perigonadal eWAT and subcutaneous iWAT) (**Fig. 5E**). Rg was unchanged in the interscapular brown adipose tissue (iBAT), heart, gastrocnemius and brain. Together, these data suggest that despite much lower insulin levels during the clamp period, multiple tissues including the liver of SNAP25^Δ3/Δ3^ mice are more responsive to insulin than SNAP25^+/+^ mice.

### Elevation of tissue norepinephrine availability and increased beiging of white adipose tissue in SNAP25^Δ3/Δ3^ mice following high fat diet challenge

We then sought to investigate the changes that may have occurred in the adipose tissue of the SNAP25^Δ3/Δ3^ mice that could enhance their response to insulin and augment glucose uptake. In response to HFD, SNAP25^+/+^ mice developed normal remodeling and adipocyte hypertrophy of iBAT and iWAT (**Fig. 6A-E**). This contrasts with SNAP25^Δ3/Δ3^ mice, where adipocyte size was significantly smaller in both iWAT and iBAT (**Fig. 6A-E**). The iBAT of HFD-fed SNAP25^+/+^ mice had many large unilocular adipocytes, whereas SNAP25^Δ3/Δ3^ had smaller multilocular adipocytes (**Fig. 6A**). F4/80 staining revealed that iWAT macrophages in SNAP25^+/+^ mice were predominately located in crown-like structures but were more evenly distributed in SNAP25^Δ3/Δ3^ mice (**Fig. S4B**). As has been observed in humans, increased beiging and the accompanying remodeling of the white adipose depots to be more oxidative may be responsible for some of the increased glucose uptake into WAT in SNAP25^Δ3/Δ3^ mice (26). Indeed, as in the chow-fed mice, HFD-fed SNAP25^Δ3/Δ3^ mice exhibited increased iWAT beiging (**Fig. 6B**). Recent studies have observed an increase in the quantity of sympathetic nerve endings in beiged adipose tissue (27–32). Consistent with this, immunohistochemical staining of adipose tissue sections for Tyrosine Hydroxylase (TH), the first and rate-limiting enzyme in the catecholamine biosynthesis pathway, showed a significant increase in the number of TH-positive sympathetic innervations in both iBAT and iWAT (quantified in **Fig. 6C,D**). This increase in TH-positive nerves in the iWAT is evident in the stained sections (**Fig. 6B,D**).

**Figure 6.**
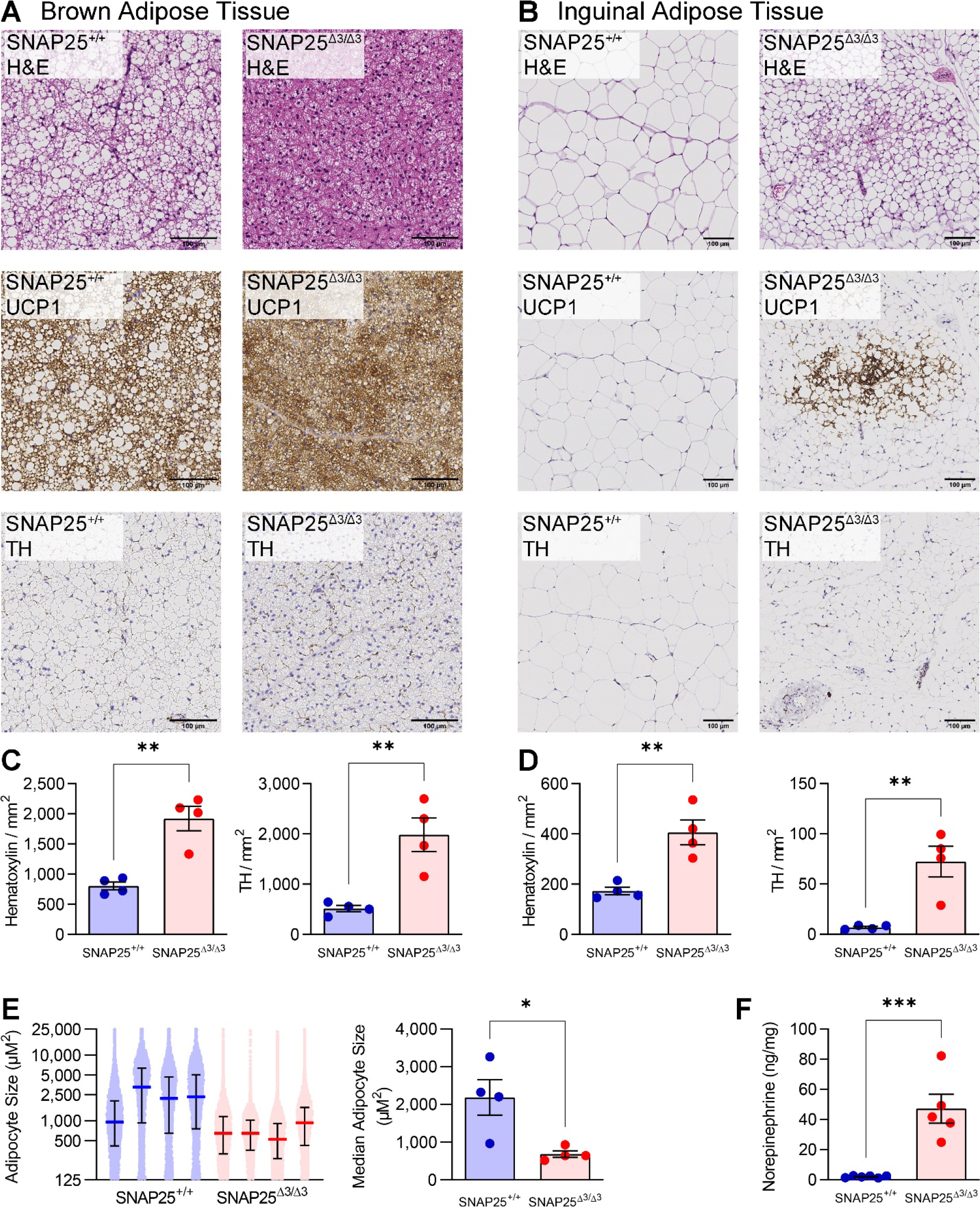
**Adipose tissue remodeling in response to HFD is replaced by increased norepinephrine-PKA signaling and beiging in SNAP25**^Δ**3/**Δ**3**^ **mice A.** Representative H&E-, UCP1-, and TH-stained sections of iBAT. **B.** Representative H&E-, UCP1-, and TH-stained sections of iWAT. **C.** Quantification of nuclei (hematoxylin) and TH-positive neurons in the iBAT TH-stained slides. **D.** Quantification of nuclei (hematoxylin) and TH-positive neurons in the iWAT TH-stained slides. Images are a representative sample from three to four mice from each group. **E.** Quantification of iWAT adipocyte size in four mice of each genotype. On the left, each dot represents the area of an individual adipocyte, and each mouse is plotted separately. Median and interquartile range are plotted. On the right, the median iWAT adipocyte size for each mouse is plotted. **F.** Norepinephrine content of iWAT from HFD-fed SNAP25^+/+^ and SNAP25^Δ3/Δ3^ male mice. *N* = 6 SNAP25^+/+^, 5 SNAP25^Δ3/Δ3^. ** p<0.01, *** p<0.001. Values are expressed as mean ± SEM. Analyses were performed using t-test.

The SNAP25Δ3 mutation prevents the Gβγ subunit from inhibiting vesicular membrane fusion and thereby the exocytosis of a currently undetermined number of neurotransmitters and hormones. There are a number of hormones that can elicit adipose tissue browning (6, 33). We sought to determine which hormones are altered as this would shed light onto the mechanism behind the metabolic alterations in SNAP25^Δ3/Δ3^ mice. The archetype signaling mechanism to induce adipocyte browning is cold stimulating the sympathetic nervous system to release the catecholamine NE into adipose tissue, which stimulates cAMP-PKA signaling events in adipocytes through the βARs (34). Because of this, we measured catecholamines by high-performance liquid chromatography (HPLC) in these HFD-fed mice. Although circulating catecholamines were unchanged (**Fig. S5**), measurements directly from homogenized iWAT (2.06 ± 0.26 vs. 47.20 ± 9.61 ng/mg; SNAP25^+/+^ vs. SNAP25^Δ3/Δ3^; mean ± SEM) revealed a 23-fold increase in NE **(Fig. 6F).** These data indicate that in SNAP25^Δ3/Δ3^ mice excess NE in the adipose tissues promotes beiging.

### Attenuation of the acute hyperphagic response to high fat diet in SNAP25^Δ3/Δ3^ mice

To determine the mechanism for the relative reduction in weight gain and fat mass in SNAP25^Δ3/Δ3^ mice, we assessed lipid absorption, food intake, and energy expenditure in mice on HFD. Fecal triglyceride content was not different (1.10 ± 0.29 vs. 0.86 ± 0.21 μg triglyceride/mg feces; *N* = 8 SNAP25^+/+^ vs. 7 SNAP25^Δ3/Δ3^; mean ± SEM; P = 0.526), indicating that the reduction in body weight in SNAP25^Δ3/Δ3^ mice was not due to a decrease in lipid absorption by the gut. We examined food intake over the first 5 weeks as chow-fed mice were switched to a HFD (**Fig. 7A**). During the first two weeks of HFD, SNAP25^+/+^ mice exhibit a robust increase in food intake, which was attenuated in SNAP25^Δ3/Δ3^ mice (**Fig. 7A**). By week 4 the daily food consumption was similar between genotypes. To assess more empirically whether the SNAP25^Δ3/Δ3^ mice might be unable to mount a robust hyperphagic response we utilized a fasting-refeeding paradigm where animals were fasted for 12 h and then presented with a HFD (**Fig. 7B**). Both genotypes had an equivalent and marked increase in HFD consumption over a 4- hour period. To assess food preference, we presented SNAP25^Δ3/Δ3^ mice and their SNAP25^+/+^ littermates with HFD and chow in tandem and monitored total intake of both diets and body weight **(Fig. 7C**). No significant differences were identified based on genotype. In summary, we observed that SNAP25^Δ3/Δ3^ mice do not overeat the more palatable, caloric dense HFD to the same extent as their SNAP25^+/+^ littermates, but this is not driven by an aversion to HFD or the inability to mount a hyperphagic response.

**Figure 7.**
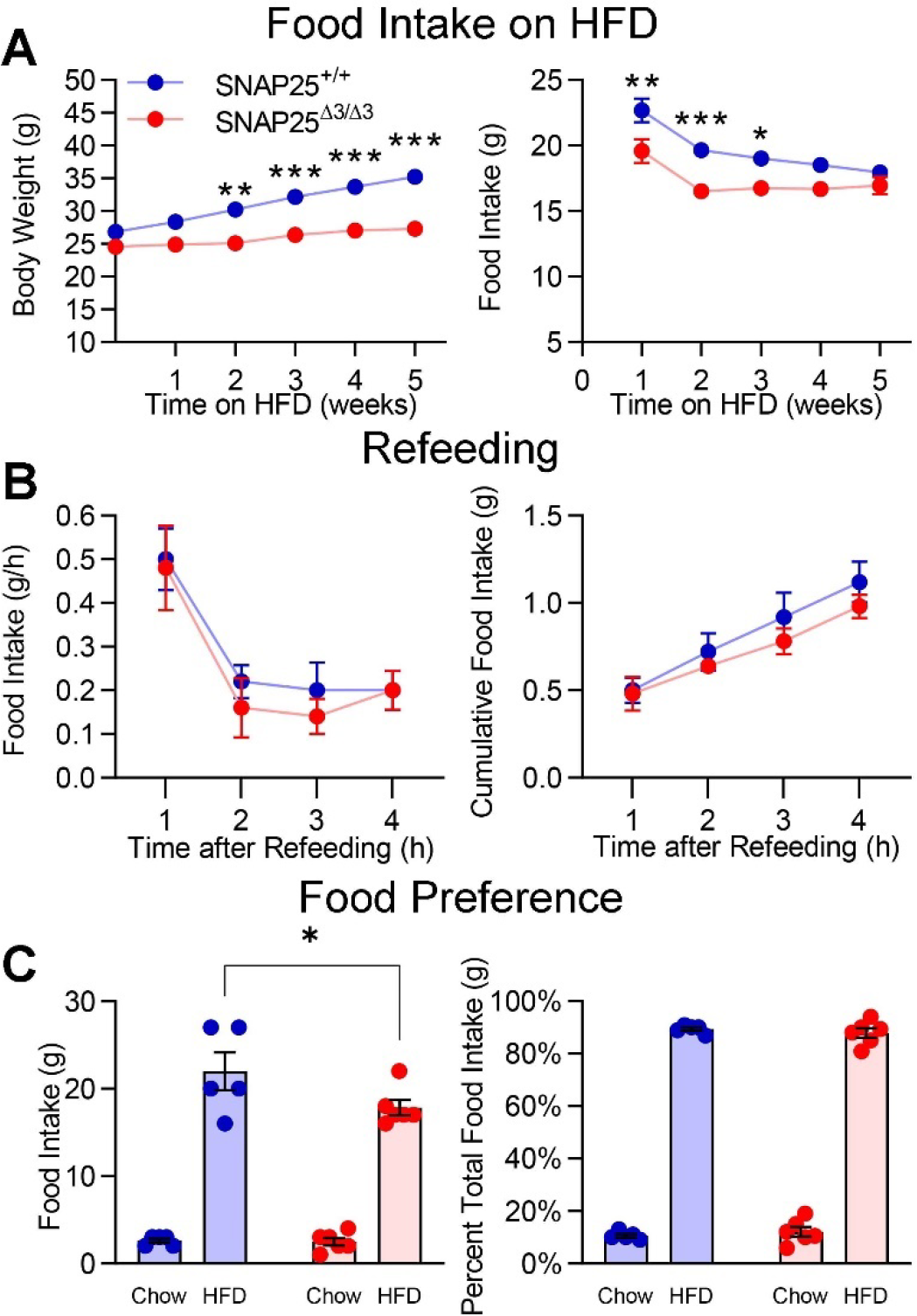
**Initial hyperphagic response to HFD is attenuated without differences in fasting- induced hyperphagia or dietary preference in SNAP25**^Δ**3/**Δ**3**^ **mice A.** Body weight (P < 0.001, effect of genotype; P < 0.001, effect of genotype × time interaction) and food consumption (P < 0.001, effect of genotype; P = 0.2466, effect of genotype × time interaction) during the first 5-weeks of HFD feeding in a separate cohort of male SNAP25^+/+^ and SNAP25^Δ3/Δ3^ mice. *N* = 23 SNAP25^+/+^, 18 SNAP25^Δ3/Δ3^. **B.** Consumption of HFD in male SNAP25^+/+^ and SNAP25^Δ3/Δ3^ mice following a 12-hour fast. *N* = 5 SNAP25^+/+^, 5 SNAP25^Δ3/Δ3^. **C.** Cumulative food intake from chow adapted male SNAP25^+/+^ and SNAP25^Δ3/Δ3^ mice given *ad libitum* access to both chow and HFD. *N* = 5 SNAP25^+/+^, 6 SNAP25^Δ3/Δ3^. * p<0.05, ** p<0.01, *** p<0.001. Values are expressed as mean ± SEM. Analyses were performed using two-way ANOVA and post hoc analyses were performed using Bonferroni multiple comparisons test for SNAP25 genotype only and are indicated on figures.

A more thorough assessment of this transition from chow to HFD was performed using the Promethion calorimetry system to continuously monitor diurnal feeding patterns and energy expenditure (**Fig. 8**). Energy expenditure was not different and was highest in the dark phase in both genotypes on a chow diet (**Fig. 8B**). When presented with a HFD both genotypes showed a robust and comparable increase in energy expenditure and caloric intake (**Fig. 8B,C**). While energy expenditure did not differ between genotypes during the 12 days on a HFD (**Fig. 8B**), the initial hyperphagia gradually subsided in the SNAP25^Δ3/Δ3^ to a greater extent than in SNAP25^+/+^ mice (**Fig. 8C**). The 12-day cumulative caloric intake (**Fig. 8A**), and consequently adiposity (**Fig. 8E**), was decreased in SNAP25^Δ3/Δ3^ mice. Thus, SNAP25^Δ3/Δ3^ mice have a short-term but significantly altered food consumption pattern, which contributes to the reduced weight gain on a HFD. Taken together we can infer that altered feeding behavior, in particular the initial response to HFD, in SNAP25^Δ3/Δ3^ mice is a contributor to the differential weight gain phenotype.

**Figure 8.**
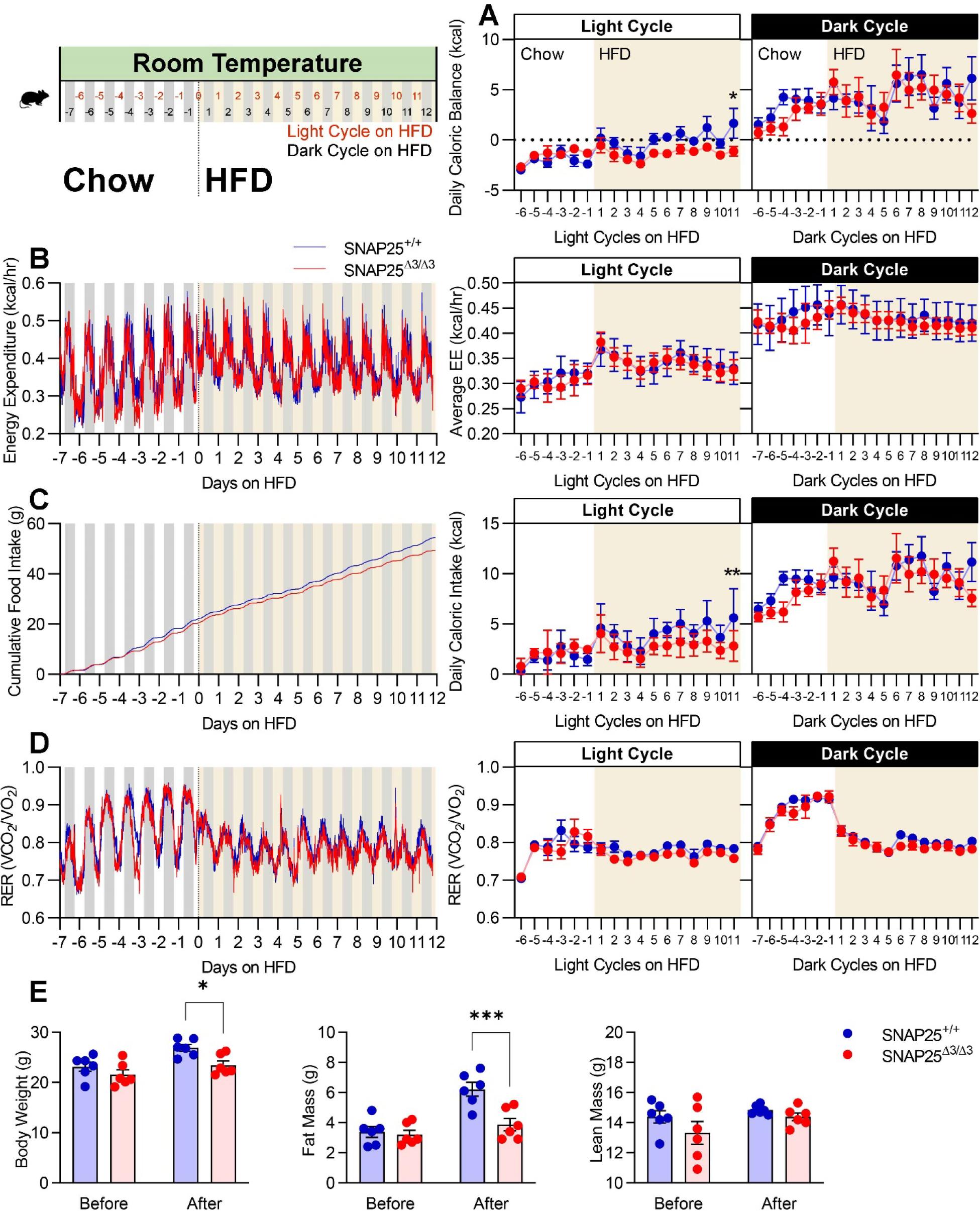
**Lower net caloric accumulation upon transition to HFD is associated with lesser food intake without alterations in energy expenditure in SNAP25**^Δ**3/**Δ**3**^ **mice**Chow-fed twelve-week-old male SNAP25^+/+^ and SNAP25^Δ3/Δ3^ mice were acclimated in the Promethion System for one week prior to transitioning to HFD. **A.** Energy balance (P = 0.0006 for the effect of genotype × time during the light cycle). **B.** Energy expenditure. **C.** Cumulative food intake (P = 0.0556 for the effect of genotype; P = 0.0023 for the effect of genotype × time during the light cycle). **D.** Respiratory exchange ratio. **E.** Body weights and composition before (day -7) and after (day 12) these energy balance studies. For all figures, *N* = 6 SNAP25^+/+^, 6 SNAP25^Δ3/Δ3^. * p<0.05, *** p<0.001. Light and dark cycles were analyzed separately. Analyses were performed by two-way ANOVA with repeated measures and post hoc analyses were performed using Bonferroni multiple comparisons test for SNAP25 genotype only. Energy expenditure was assessed using ANCOVA.

### Reversal of metabolic protection in SNAP25^Δ3/Δ3^ mice by thermoneutral housing conditions

A mild thermogenic stress is evoked in mice by housing at standard room temperature (∼22 °C). This requires activation of the autonomic nervous system in adipose and other tissues to support thermoregulation (37). Increased autonomic drive and beiging of adipose tissue are observed in the SNAP25^Δ3/Δ3^ mice housed at standard housing conditions (**Fig. 6**), suggesting that the Gβγ-SNARE interaction is a negative regulator of autonomic activity. We hypothesized that if we minimized the induction of autonomic drive by shifting animals to a thermoneutral setting (∼30 °C) the metabolic benefits of SNAP25^Δ3/Δ3^ would decrease. Therefore, we housed SNAP25^Δ3/Δ3^ mice and their SNAP25^+/+^ littermates in thermoneutral housing for four weeks prior to providing them a HFD. Indeed, the reduced weight gain, adiposity, and fat pad size of SNAP25^Δ3/Δ3^ mice in response to a HFD were eliminated (**Fig. 9A-C**). Food intake in response to HFD feeding was also no longer different between genotypes during the thermoneutral housing conditions (**Fig. 9D**). Thermoneutral housing did not revert SNAP25^Δ3/Δ3^ food intake to SNAP25^+/+^ room temperature levels; instead, in both genotypes food intake was lower than in room temperature housed animals during all weeks of the study (**Fig. 7A, 9D**). In addition, the improvements in insulin action of SNAP25^Δ3/Δ3^ mice during IP-GTT at room temperature, in both the chow and HFD feeding period, were also abolished by housing the mice in thermoneutral conditions (**Fig. 9E,F**). These findings demonstrate that the metabolic improvements caused by removing the Gβγ-SNARE interaction are dependent upon the environmental temperature in which the mice are housed. This is consistent with our hypothesis that increased sympathetic activity due to loss of the α -adrenergic brake to exocytosis mediates the metabolic protection conferred in SNAP25^Δ3/Δ3^ mice.

**Figure 9.**
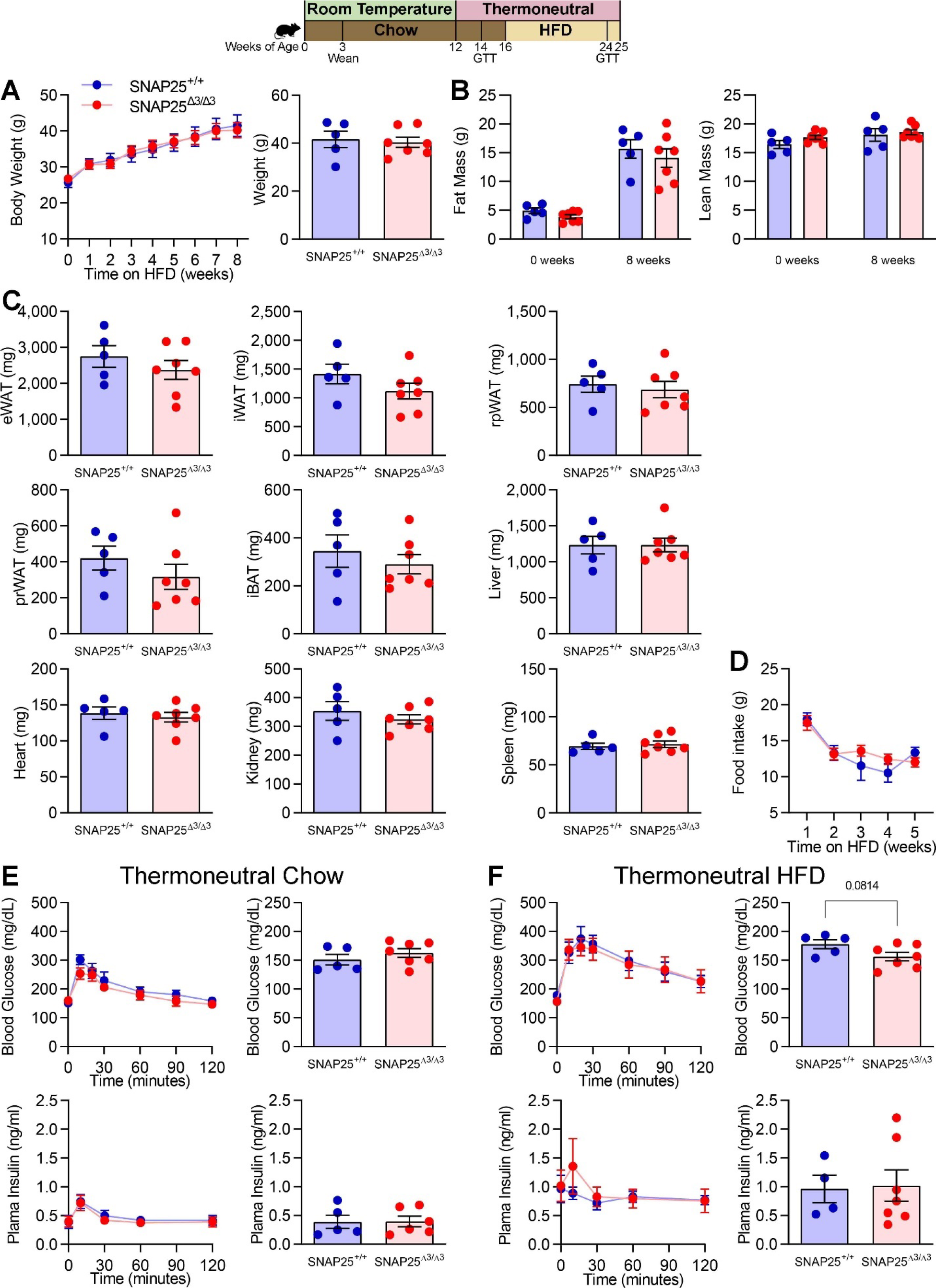
**Lack of metabolic protection during HFD feeding in SNAP25**^Δ**3/**Δ**3**^ **mice under thermoneutral housing conditions A.** Following 4 weeks of acclimation to thermoneutral housing conditions, male SNAP25^+/+^ and SNAP25^Δ3/Δ3^ were fed HFD for 8 weeks. Comparison of terminal body weights. **B.** Body composition at beginning and end of HFD feeding period. **C.** Tissue weights of SNAP25^+/+^ and SNAP25^Δ3/Δ3^ mice at the end of the HFD feeding period. **D.** Food consumption during the first 5-weeks of HFD feeding during thermoneutral housing conditions. *N* = 5 SNAP25^+/+^, 7 SNAP25^Δ3/Δ3^. Analysis was performed by two- way ANOVA with repeated measures and post hoc analyses were performed using Bonferroni multiple comparisons test for SNAP25 genotype only. **E.** Blood glucose and plasma insulin levels during an IP- GTT in chow-fed male SNAP25^+/+^ and SNAP25^Δ3/Δ3^ housed at thermoneutral conditions for two weeks. Fasting blood glucose and plasma insulin at “time 0” of GTT. **F.** Blood glucose and plasma insulin levels during an IP-GTT in HFD-fed male SNAP25^+/+^ and SNAP25^Δ3/Δ3^ housed at thermoneutral conditions. Fasting blood glucose and plasma insulin at “time 0” of GTT. For all figures *N* = 5 SNAP25^+/+^, 6-7 SNAP25^Δ3/Δ3^. Values are expressed as mean ± SEM. Most analyses were performed by two-way ANOVAs with repeated measures and post hoc analyses were performed using Bonferroni multiple comparisons test for SNAP25 genotype only. Exceptions were terminal body weight (A right), tissue weights (C), and fasting glucose and insulin (D and E right), which were performed using t-test.

### Prevention of diet induced remodeling of adipose tissues in SNAP25^Δ3/Δ3^ mice by thermoneutral housing conditions

Increasing sympathetic activity to induce heat production from adipose tissue is a fundamental part of the response to environmental cold stress (38, 39). In the mice housed at thermoneutral conditions, this adipose beiging and elevation of NE content is eliminated (**Fig. 10A-F**). Both iBAT and iWAT from SNAP25^Δ3/Δ3^ mice are remodeled as expected in response to HFD feeding (**Fig. 10A-E**), but no difference is seen between the genotypes. In both genotypes, macrophages in iWAT are predominately located in crown-like structures (**Fig. S4B**). The ability of thermoneutral housing conditions to reverse the metabolic improvements achieved by disrupting the Gβγ-SNARE interaction indicates that these improvements are related to an elevation of autonomic activity in tissues such as adipose tissue.

**Figure 10.**
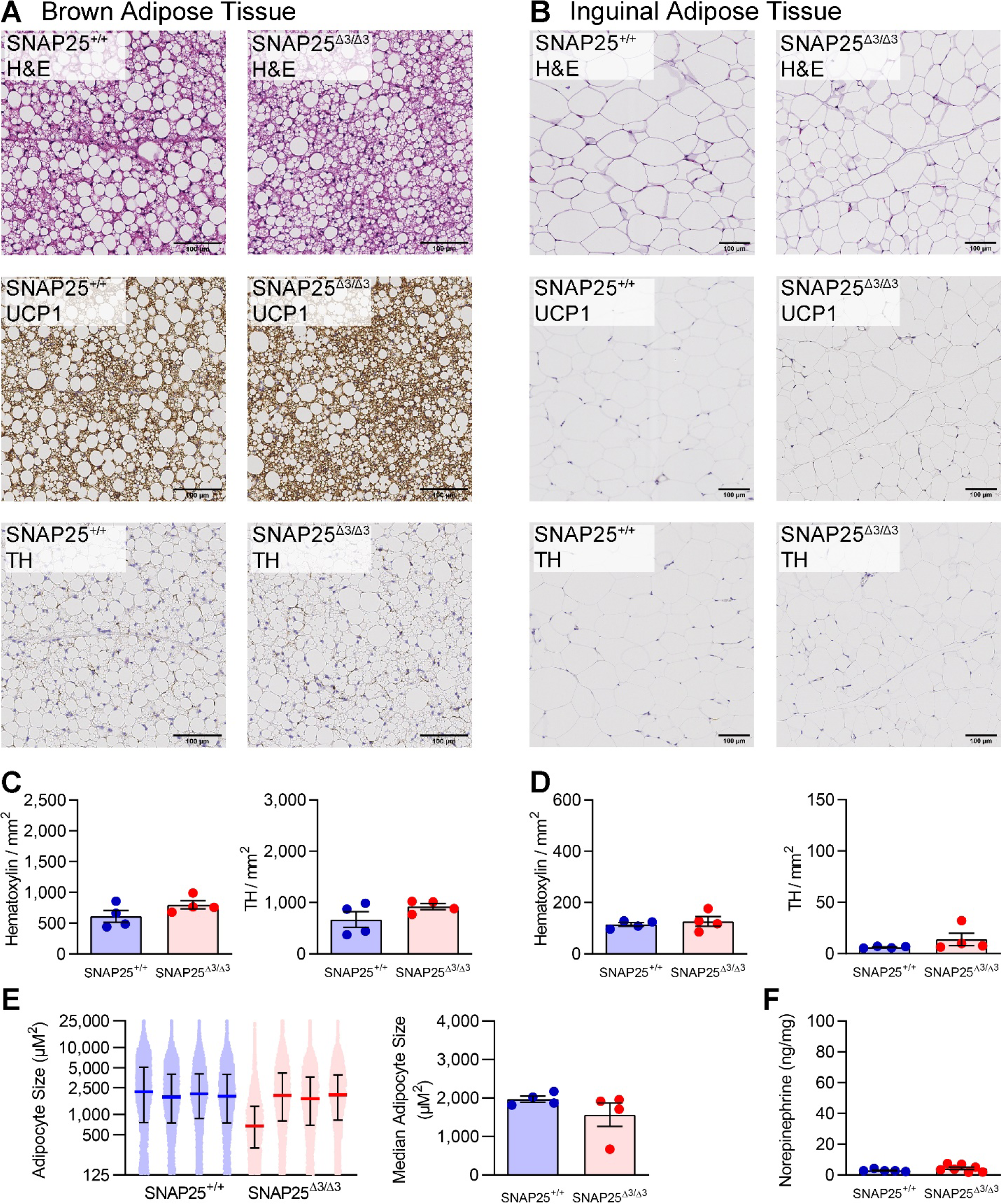
**Adipose tissue remodeling in response to HFD is not different in SNAP25**^Δ**3/**Δ**3**^ **mice during thermoneutral housing conditions A.** Representative H&E-, UCP1-, and TH-stained sections of iBAT. **B.** Representative H&E-, UCP1-, and TH-stained sections of iWAT. **C.** Quantification of nuclei (hematoxylin) and TH-positive neurons in the iBAT TH-stained slides. **D.** Quantification of nuclei (hematoxylin) and TH-positive neurons in the iWAT TH-stained slides. Images are a representative sample from three to four mice from each group. **E.** Quantification of iWAT adipocyte size in four mice of each genotype. On the left, each dot represents the area of an individual adipocyte, and each mouse is plotted separately. Median and interquartile range are plotted. On the right, the median iWAT adipocyte size for each mouse is plotted. **F**. Norepinephrine content of iWAT from HFD-fed SNAP25^+/+^ and SNAP25^Δ3/Δ3^ male mice. *N* = 5 SNAP25^+/+^, 7 SNAP25^Δ3/Δ3^. Values are expressed as mean ± SEM. All analyses were performed using t-test.

### Release of norepinephrine from sympathetic endings is amplified in SNAP25^Δ3/Δ3^ mice

To directly test the hypothesis that removing Gβγ inhibition of SNARE-mediated vesicular fusion causes more NE to be secreted when sympathetic neurons are excited, we measured NE release from electrode-stimulated sympathetic nerves innervating iWAT using a selective genetically encoded NE biosensor, GRAB_NE_ (40). This allows for good spatiotemporal resolution of junctional NE secretion. GRAB_NE_ expression was placed under the control of *TH*-Cre, which limits its expression to catecholaminergic and dopaminergic neurons. Individual neuronal processes adjacent to adipocytes from excised iWAT were imaged using lattice light sheet fluorescence microscopy (**Fig. S6A,B**) (41–43). When exogenous NE was applied to sympathetic neurons innervating iWAT, fluorescence was stimulated in GRAB_NE_ expressing neurons (**Fig. S6C**). Plotting the fluorescence over time showed that the GRAB_NE_ was rapidly excited by NE (**Fig. S6D**). High concentrations of epinephrine stimulated GRAB_NE_, but to a much lesser extent demonstrating the fidelity of GRAB_NE_ for NE (**Fig. S6E**). Electrode-stimulation of sympathetic nerves with a train of stimuli led to repeatable transient fluorescent puncta on the sympathetic nerve endings (**Fig. 11A**). GRAB_NE_ fluorescence was localized to the rim adjacent to the round cell bodies of adipocytes.

**Figure 11.**
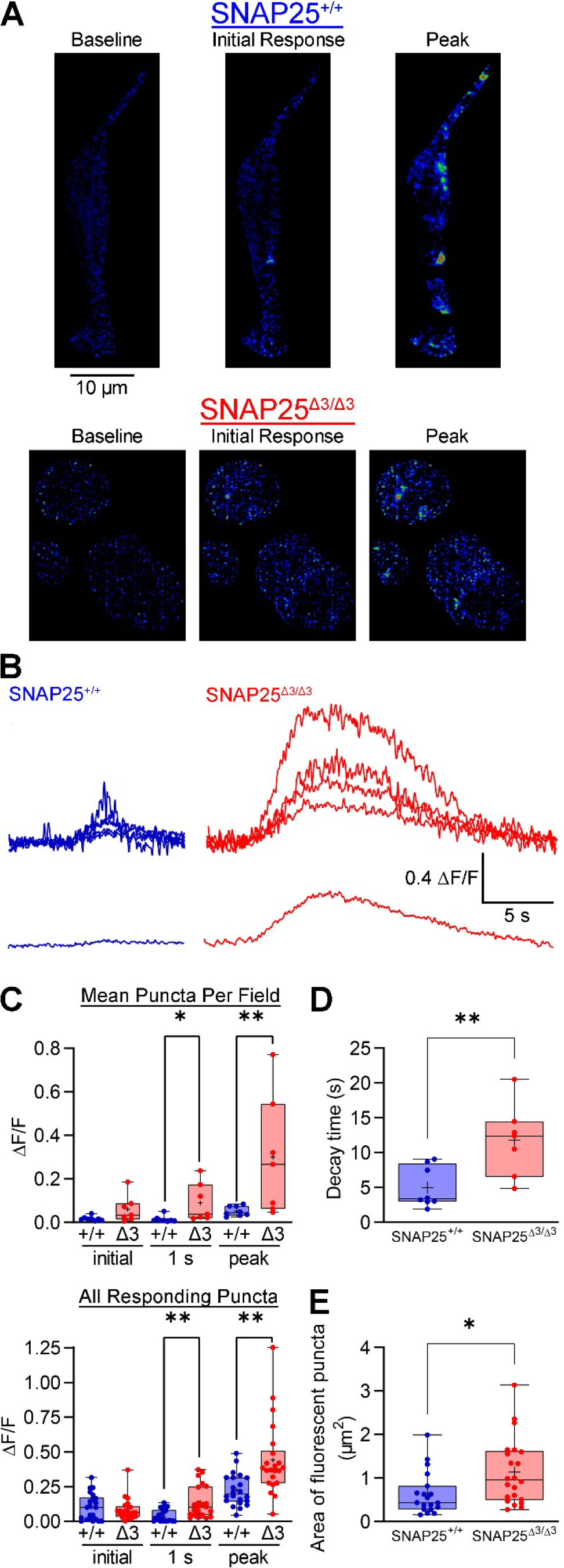
Norepinephrine release measured *ex vivo* from sympathetic neurons in the inguinal adipose depot is greater in SNAP25^Δ**3/**Δ**3**^ mice. **A.** Live cell images of 535nm GRAB_NE_ fluorescence from mice hemizygous for Rosa26-LSL-GRAB-NE and *TH*-Cre showing fluorescent puncta surrounding, but not within, adipocytes before, during, and after stimulus trains of 100 stimuli, 100-200 µA at 20 Hz and 1 ms duration. Areas of low fluorescence are depicted in blue, while areas of high fluorescence are depicted in red in the false-color heatmap. **B.** Representative fluorescence transients graphed as ΔF/F from iWAT SNAP25^+/+^ (blue) and SNAP25^Δ3/Δ3^ littermates (red). **C.** Top: Bar graphs of mean ΔF/F values per field for iWAT taken from SNAP25^+/+^ and SNAP25^Δ3/Δ3^ immediately upon stimulation, 1 s post stimulus, and at the peak of the stimulus. Bottom: Bar graphs of individual ΔF/F values per responding puncta for iWAT of SNAP25^+/+^ and SNAP25^Δ3/Δ3^ immediately upon stimulation, 1 s post stimulus, and at the peak of the stimulus. **D.** Decay times in seconds from peak to baseline per field. **E.** Bar graphs of peak area responding puncta in µm^2^. *N* = 5 SNAP25^+/+^, 3 SNAP25^Δ3/Δ3^ biological replicates with all puncta displayed on graphs. * p<0.05, ** p<0.01. Values are displayed as a box-and-whisker plot with the mean being represented by “+”. All analyses were performed using t-test.

We used this system to compare the NE release profile between SNAP25^+/+^ and SNAP25^Δ3/Δ3^ iWAT sympathetic neurons (**Fig. 11B**). The initial stimulation of the sympathetic neuron axons produced a similar response (**Fig. 11C**). At both one second after stimulation and during the maximal response, the stimulus evoked GRAB fluorescence in sympathetic neurons from SNAP25^Δ3/Δ3^ mice was significantly greater (**Fig. 11C**). The time required for fluorescence to return to baseline was also greater in SNAP25^Δ3/Δ3^ (**Fig. 11D**). Additionally, SNAP25^Δ3/Δ3^ mice had a greater area of fluorescence response consistent with the idea that these mice have more iWAT sympathetic innervation (**Fig. 11E**). These findings are consistent with our prediction that the SNAP25Δ3 mutation removes a brake on NE exocytosis without affecting the initial exocytosis event. In summary, in this experiment we used electrical stimulation to directly provoke NE secretion from sympathetic nerves in iWAT. This allowed us to specifically measure the NE release profile from iWAT sympathetic nerves, which in SNAP25^Δ3/Δ3^ mice we found to be increased in both amplitude and duration. This is the likely mechanism by which mild cold-stimulation of the sympathetic nervous system caused by room temperature housing leads to the large increase in iWAT NE content and UCP1-positive adipocytes in SNAP25^Δ3/Δ3^ mice.

We posit that the fundamental process driving the phenotype of this mouse lacking the Gβγ brake on SNARE-mediated vesicle fusion is amplified neurotransmitter release from the sympathetic nervous system. This was most apparent in the exacerbated sympathetic response to the mild cold stress caused by housing mice at room temperature, which enhanced the release of NE leading to the induction of the adipocyte thermogenic program (**Fig. 12**). This increased sympathetic activity likely affected other metabolically active organs. NE induction of adipose tissue beiging and the thermogenic program in SNAP25^Δ3/Δ3^ mice likely contribute to improvements in insulin action particularly in the adipose tissue. This increased sympathetic neural drive causes SNAP25^Δ3/Δ3^ mice, which only lack the inhibitory actions of Gβγ on the SNARE complex, to be remarkably resistant to diet induced obesity and the associated impairment in glycemic control.

**Figure 12.**
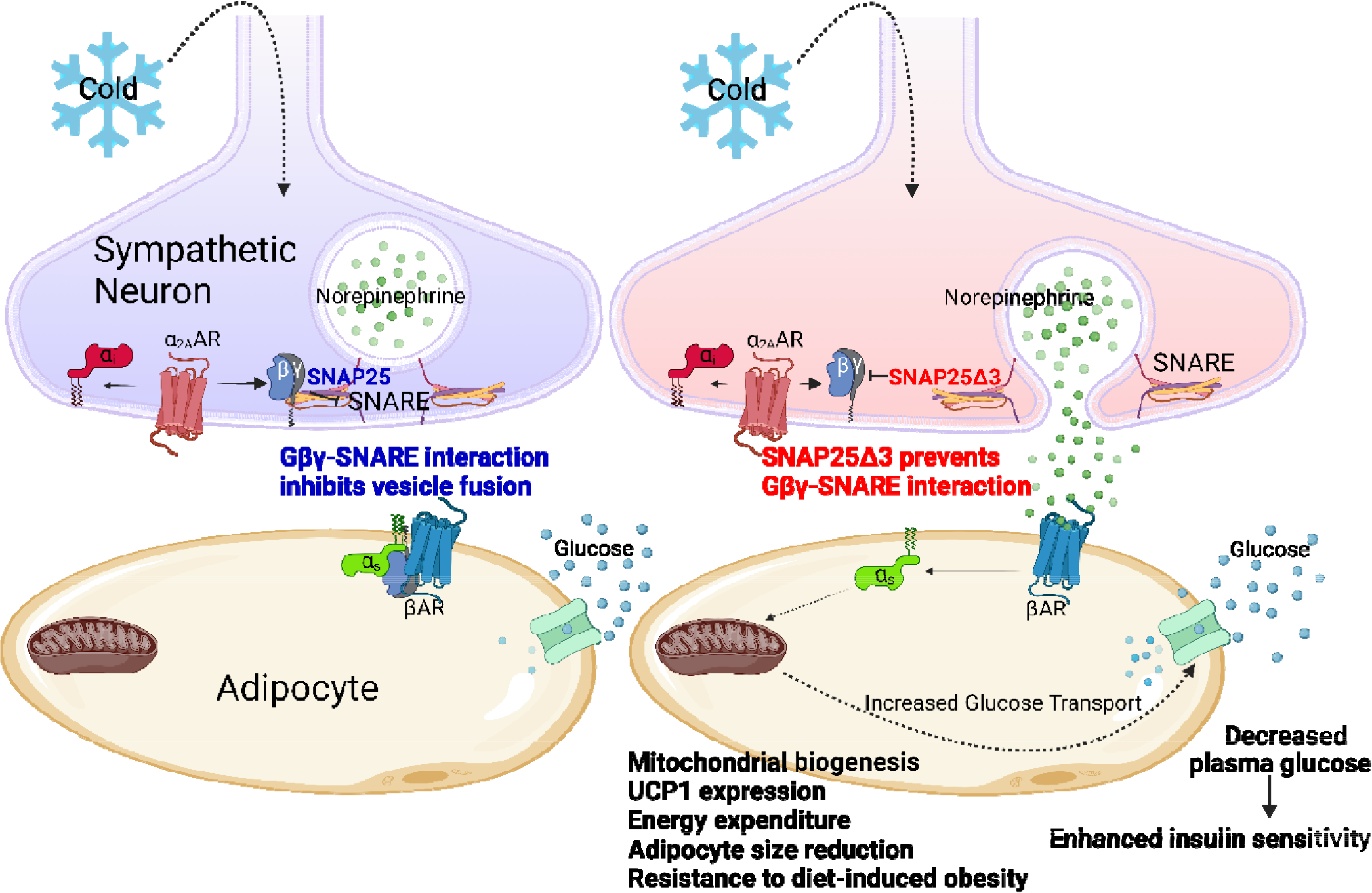
Summary. During cold stress, sympathetic regulation of adipocyte beiging/browning occurs through release of NE from sympathetic neurons and thereby induction of the adipocyte β-adrenergic program. This increases mitochondrial biogenesis (browning) and expression of UCP1 generating heat by expending energy through uncoupled respiration. Housing mice at room temperature (normal housing is around 22 °C) evokes a mild cold stress that is generally unnoticed. Normally there is an α_2A_-adrenergic brake on neurotransmitter vesicle fusion that works through the Gβγ-SNARE interaction limiting NE release. We have removed this brake on NE release by creating a mutant SNAP25 that decreases Gβγ binding leading to enhanced and prolonged NE release from sympathetic neurons and thereby greater induction of the adipocyte thermogenic program. This enhanced browning increases glucose uptake into beige adipocytes leading to enhanced insulin sensitivity, particularly in these white adipose depots. We posit that this global genetic alteration enhances and prolongs synaptic vesicle exocytosis from SNAP25 expressing neurons associated other organ systems, such as those responsible for food intake. A result of these adaptations, SNAP25^Δ3/Δ3^ mice are resistance to diet-induced obesity. Removal of the cold-induced sympathetic activity by placing animals under thermoneutral conditions (∼30 °C) reverses all of these changes, allowing us to conclude that the inability of Gβγ to act as a negative regulator of NE release is the main driver of the beneficial metabolic phenotype.

## Discussion

We began these studies to examine the role of the Gβγ-SNARE interaction in the fusion and secretion of insulin containing vesicles. Though SNAP25 is primarily expressed in neural populations (45, 46), it is also highly expressed in the β-cells of pancreatic islets (47, 48). Prior studies using an immortalized β-cell line showed that inhibition of insulin secretion by the α_2_AR autoreceptor occurs in part through Gβγ SNARE interaction (13). Our SNAP25^Δ3/Δ3^ mice provided the first opportunity to test these findings both *ex vivo*, using intact islets, and *in vivo*, by measuring circulating insulin in response to a glucose challenge. However, we found no evidence that this negative regulation of insulin secretion by α AR was removed in SNAP25^Δ3/Δ3^ mice, though this could be explained by differences between isolated *ex vivo* islets and the cultured β-cell line. Another possibility is that in islet β-cells a protein other than SNAP25 provides the scaffolding for the Gβγ interaction with insulin granules (49–51), such as SNAP23 (52). During the course of these studies, we found that lean, chow-fed SNAP25^Δ3/Δ3^ mice did not secrete as much insulin, though glucose levels were similar during GTTs. This indicates that insulin dependent glucose disposal was enhanced. Studies utilizing glucose clamps confirmed that insulin action is improved in SNAP25^Δ3/Δ3^ mice. This compelled us to conduct a series of studies to determine why these SNAP25^Δ3/Δ3^ mice would have improved insulin sensitivity.

Our first clue to the mechanism behind these phenotypes came from the observation that SNAP25^Δ3/Δ3^ mice have increased beiging of their iWAT. This was particularly noticeable in SNAP25^Δ3/Δ3^ mice challenged with HFD, since the histology revealed extensive patches of beige adipocytes in iWAT. This partially explains the improved insulin sensitivity observed in SNAP25^Δ3/Δ3^ mice as the glucose tracer studies showed increased uptake in iWAT. We did not see any notable increase in energy expenditure due to the adipocyte beiging in SNAP25^Δ3/Δ3^ mice. Therefore, the real benefit of beiging in this context appears to be enhancing insulin action on adipose tissue and other tissues such as liver and muscle (53, 54). Though iWAT beiging may not account for the largest proportion of the change in glucose uptake in SNAP25^Δ3/Δ3^ mice, it is a notable contributor to the improvement in insulin action and is very striking.

A substantial number of hormones are known to influence the adipocyte thermogenic program, many of them through GPCR-mediated signaling events, but these are mediated by the α-subunits and their effects on cAMP-PKA signaling pathways (6). This, and reports that SNAP25 is not expressed in adipocytes (55–58), led us to suspect that the iWAT beiging phenotype was not due to the actions of Gβγ-SNARE in the adipocytes themselves, but rather some hormone that is dysregulated in SNAP25^Δ3/Δ3^ mice. We initially suspected sympathetic signaling because SNAP25^Δ3/Δ3^ mice have an enhanced thermogenic response to handling stress (16, 25), but this was discounted after we failed to observe a differential response with 6 °C cold challenge or in circulating catecholamine levels.

We sought to come up with a way to effectively “turn off” the actions of the SNAP25Δ3 mutation. Our prior work shows that SNAP25^Δ3/Δ3^ mice have an exacerbated response to stress (16, 25). It is known that room temperature housing is a mild cold stress for mice (59, 60) and that cold stress can induce WAT beiging (61). So, we hypothesized that a heightened response to even to moderate cold stress could account for some of the phenotypic differences observed in SNAP25^Δ3/Δ3^ mice. When housed under thermoneutral conditions the metabolic phenotype of SNAP25^Δ3/Δ3^ mice was completely normalized. This strongly indicated that sympathetic activity is altered in room temperature housed SNAP25^Δ3/Δ3^ mice, so we revisited the idea that NE levels might be different measuring it in frozen iWAT samples. We found an enormous 23-fold increase in NE in the room temperature housed mice, which was completely returned to baseline in samples from mice housed under thermoneutral conditions. Because all metabolic differences were normalized by thermoneutral housing, and these differences cannot be accounted for solely by the lack of iWAT beiging, we posit that autonomic activity is increased in room temperature housed SNAP25^Δ3/Δ3^ mice driving the metabolic protection.

We can infer from the studies discussed thus far that in iWAT, removing this Gβγ brake amplifies the cold-stimulated NE secretion. We previously showed that α_2A_ARs work through Gβγ- SNARE interactions (16). To directly test this mechanism, we used the GRAB_NE_ fluorescent NE biosensor to unambiguously measure the NE release profile from the iWAT sympathetic neurons of these mice (40). Direct stimulation of sympathetic nerves in iWAT in SNAP25^Δ3/Δ3^ mice does not affect the initial NE exocytosis but rather intensifies and prolongs the NE release. The effect of SNAP25 genotype upon the timing of the NE release profile is consistent with a model of defective α_2A_AR autoreceptor signaling, where junctional NE must first rise to exert negative feedback. We posit that loss of the brake to SNARE-mediated vesicle fusion and neurotransmitter release enhances the net release of NE leading to increased autonomic activity.

The metabolic phenotype of the SNAP25^Δ3/Δ3^ mouse is more complex than just WAT beiging.

The altered feeding behavior of SNAP25^Δ3/Δ3^ mice appears to be a major contributor to their resistance to diet induced obesity. Upon calculation of net energy balance (total energy expenditure-caloric intake), SNAP25^+/+^ mice tended to remain in a more positive energy balance than SNAP25^Δ3/Δ3^ mice during HFD feeding. This initial relative hypophagia in SNAP25^Δ3/Δ3^ mice is likely a major contributor to their resistance to HFD-induced obesity as we did not observe any differences in fasting-induced hyperphagia or food preference. The fact that differences in food intake are reversed at thermoneutrality suggests that this is also controlled by autonomic activity. This finding might be predicted since environmental temperature and the sympathetic nervous system are known to affect feeding behavior (62–64). Given the impact of central autonomic drive for modulating food intake and energy expenditure, future studies could target specific brain regions to integrate this Gβγ-SNARE mechanism into our current understanding of the complex circuitry controlling appetite.

In summary, we have found that disabling the Gβγ-SNARE brake on exocytosis in mice *in vivo* causes an exacerbation of the mild cold stress from room temperature housing, leading to elevated autonomic NE release. Our findings parallel those of a recent study using a brain-sparing sympathofacilitator (PEGylated amphetamine) to increase autonomic activity, which saw increases in WAT NE levels and resistance to diet induced obesity occur (65). Retrospectively, it makes sense that amphetamines, which inhibit the reuptake of monoamine neurotransmitters, and the SNAP25^Δ3/Δ3^ mouse would have phenotypic similarities, such as reduced food intake, increased iWAT beiging, and resistance to diet induced obesity. These studies highlight the importance of considering changes in the sympathetic nervous system when evaluating metabolic phenotypes as even slight alterations in autonomic activity can have profound physiological consequences. Our findings also demonstrate how the chronic thermal stress due to housing mice at room temperature can have a drastic effect on phenotype. These studies present a striking example of an interaction between genotype and thermal environment. Mechanistically, we clearly show that disabling the Gβγ-SNARE interaction in sympathetic neurons innervating iWAT enhances and prolongs NE release upon stimulation. Pharmacological targeting of this Gβγ-SNARE interaction could be an interesting alternative to directly targeting the GPCRs themselves, given the broad improvements in food intake, insulin action, and adipose browning, which all occurred without cardiovascular effects. Any potential neurological side effects could be ameliorated through designing CNS sparing agents, while avoiding GPCR targets that could have broad tissue distributions and side effects. Thus, the approach of targeting a pathway that modulates rather than drives specific neuroendocrine signals is a novel approach to treat metabolic disease.

## Materials and Methods

### Animal procedures

Mice used for these experiments were generated from heterozygous breeding of SNAP25^+/Δ3^ mice on a C57BL/6 background. The strategy for the generation of these mice was previously described (16). Mice were maintained on a 12-hour light, 12-hour dark cycle and housed with 3–5 animals per cage except during monitoring of feed intake when they were singly housed. Mice were housed at standard housing room temperature (∼22 °C) or thermoneutral housing conditions (∼29.5 °C) when indicated. Mice were maintained *ad libitum* on chow (13.5% calories from fat: 5001; LabDiet) or high fat diet (HFD) (60% calories from fat: 3282; Bio-Serv) when indicated. Mice were weighed when indicated and body composition was measured using a LF50 Body Composition Analyzer (Bruker) at Vanderbilt Mouse Metabolic Phenotyping Center. All studies were performed with littermates of the appropriate genotypes. Mice were fasted 5 h and euthanized by isoflurane overdose or CO_2_ asphyxiation and exsanguination via cardiac puncture at the end of the study for collection of blood and tissues. Data from one mouse was excluded from all studies due to hydronephrosis. All procedures were approved by the Institutional Animal Care and Use Committee at Vanderbilt University.

### Histology

Adipose tissues were fixed in 10% formaldehyde overnight and subsequently stored in 70% ethanol then processed routinely, embedded, sectioned and stained with hematoxylin and eosin (H&E) or immunohistochemically stained for Tyrosine Hydroxylase (ab152, Millipore Sigma) or UCP1 (ab10983, Abcam). Histology was performed by the Vanderbilt Translational Pathology Shared Resource. Whole slides were imaged at 20× with a Leica SCN400 Slide Scanner by the Digital Histology Shared Resource at Vanderbilt University Medical Center.

Quantification of TH staining was performed in QuPath version 0.3.0. One whole section of iWAT and iBAT were quantified for each mouse. Blood vessels, lymph nodes, and dissection artifacts were manually excluded from analysis. For all slides, hematoxylin stain was set at 0.63 0.682 0.371 and TH stain was set at 0.479 0.618 0.623. Counting was performed using the Fast Cell Counts (Brightfield) feature. Cell detection threshold was set at 0.3 and DAB threshold was set at 0.6. All other parameters were the default values.

Quantification of iWAT adipocyte cell size was performed in Fiji adapted from a method developed by Dr. Joseph Roland (https://www.vumc.org/dhsr/sites/default/files/public_files/Fiji-Adipose-Segmentation.pdf). Briefly, a TIFF image of the entire iWAT section at a resolution of 0.5 µm per pixel was uploaded into Fiji and the area outside of the tissue was manually cropped out. The image was converted to 8-bit and colors were inverted. Background was subtracted using the settings of Rolling Ball Radius 20.0 pixels and Sliding Paraboloid. The image was manually cropped into multiple segments to reduce file size. Morphological Segmentation with Gaussian radius 3 and Watershed Segmentation tolerance 4 was run on each segment. An image was created showing the dams from the Morphological Segmentation and a Gaussian blur with a Sigma of 2.0 was added. The resulting image was converted to 8-bit and the threshold settings were manually adjusted so that the cell walls are white on a black background. Analyze Particles was run with the size range of 500-inifity producing a list of adipocyte areas in pixels. Area was converted to µm^2^ and measurements below 125 µm^2^ and above 25,000 µm^2^ were excluded. All adipocytes size measurements within this range were plotted for each individual mouse, having between 25,463 and 65,480 adipocyte measurements per mouse, to determine the median adipocyte size. Analyses comparing genotypes were performed using the median adipocyte size.

### Leptin

At the end of the chow or HFD feeding studies, mice were euthanized as described and blood was obtained postmortem by terminal cardiac puncture and placed into a microcentrifuge tube coated with EDTA. Samples were kept on ice until centrifugation to obtain plasma which was stored at -80 °C. Leptin was measured from this plasma with a Luminex 100 system (Luminex Corp) at the Vanderbilt University Hormone Assay and Analytical Services Core.

### Glucose tolerance test

For all glucose tolerance tests (GTT), mice were fasted for 5 h, and fasting blood glucose was measured from a drop of tail vein blood (Accu-Chek Aviva glucometer) at the indicated time points. GTTs in **Fig. 9** were conducted in a 29.5 °C room while all other GTTs were conducted at standard room temperature. For **Fig. 2**, glucose was given by either an intraperitoneal (IP) injection or oral gavage of 2.0 g/kg glucose. For **Figs. 5 & 9**, mice were given IP injections of 1.0 g/kg dextrose (Agri Laboratories, Ltd.) in 0.9% saline (Hospira, Inc.). Insulin concentrations during GTTs (**Figs. 2,9**) were analyzed by RIA by the Vanderbilt University Hormone Assay and Analytical Services Core. In **Fig. 5B**, mice were fasted for 5 h and blood glucose was collected from the tail vein after which the mice were subsequently euthanized by CO_2_ and EDTA plasma was collected by cardiac puncture as described, which was stored at -80 °C until insulin was measured by ELISA (Mercodia).

### Mouse islet perifusion

Pancreatic islets were isolated and perifusion assays were performed on fresh islets at the Vanderbilt Islet Procurement and Analysis Core as previously described (66). Islet preparations were equilibrated and stable baseline response established at 5.6 mmol/L glucose, and insulin secretion was stimulated with 16.7 mmol/L glucose. A dose-response curve for the inhibition of GSIS by the α_2_AR selective agonist, brimonidine (Br), was generated. Islets were matched for size and number, and assessed as islet equivalents (IEQ) (67).

### Energy balance

Food intake and energy expenditure were monitored on mice in a Promethion system (Sable Systems International) by the Vanderbilt Mouse Metabolic Phenotyping Center. The system allows for the continuous high time resolution monitoring of oxygen consumption, carbon dioxide production, feeding and activity patterns and caloric intake. Animals were housed over multiple days to acclimate to the facility. Body weight and composition was monitored before and after a calorimetry study. The calorimetry system is housed in a temperature-controlled cabinet. In some studies animals were housed at standard housing temperature (22 °C) then transitioned to a cold stress (6.0 °C). In other studies, 17-19 weeks old chow-fed mice were transitioned to HFD.

### Cardiovascular imaging

Cardiac parameters were measured by parasternal M-mode echocardiography at the VUMC Cardiovascular Physiology Core as described (68, 69). Briefly, mice had their chest fur removed with a topical depilatory agent. Ultrasound coupling gel heated to 34 °C was applied to the chest area and a linear array transducer (18-23 MHz) was positioned to obtain two-dimensional B-mode parasternal long and short axis views at the mid-ventricular level (Vevo 2100, VisualSonics). One-dimensional M-mode images were obtained for measurement in the short axis view to obtain cardiac wall thickness and chamber dimensions. Left ventricular (LV) chamber size and wall thickness were measured off-line in the M-mode from at least three consecutive beats and averaged. LV wall thickness: intraventricular septum (IVS) and posterior wall (PW) at systole and diastole; and LV internal dimensions (LVID) during systole and diastole were measured.

### Hyperinsulinemic euglycemic clamps

Clamp studies were done chronically catheterized (carotid artery and jugular vein) conscious mice (70–72). Catheters were inserted 4-5 days prior to a study by the Vanderbilt Mouse Metabolic Phenotyping Center. [3-^3^H] glucose was used to measure whole body basal and clamp glucose flux. A 4 mU·kg- 1·min-1 insulin infusion was initiated to increase insulin to a physiologic range. Red blood cells from a donor animal were infused at a constant rate to replace blood taken during study. Basal and clamp period blood samples for glucose, insulin and tracer. At the end of the clamp period, multiple tissues were collected to measure the accumulation of 2[1-^14^C]Deoxy-D-glucose(DG). Using tracer methods [3- ^3^H]-glucose and 2[1-^14^C]DG during the clamp, we assessed tissue glucose uptake and whole body (and hepatic) glucose flux.

### Catecholamines

For plasma catecholamines, in order to minimize stress-induced catecholamine release mice were cannulated and allowed to move about freely while blood was collected using a blunt needle. Canulations and blood collections were performed by the Vanderbilt Brain Institute Neurochemistry Core. For catecholamines in adipose tissue, whole flash-frozen iWAT was ground into a powder in liquid nitrogen. Powdered iWAT was poured into a tube and weighed. PCA solution (5 mM glutathione in 0.4 N perchloric acid) was added to frozen iWAT powder at a ratio of 1 ml PCA per 100 mg iWAT. Samples were homogenized then centrifuged. The clear supernatant was removed and stored at -80 °C. Catecholamines in both plasma and frozen tissue lysates were analyzed by LC/MS and HPLC respectively with electrochemical detection as previously described (78) at the Vanderbilt Hormone Assay and Analytical Services Core.

### Fecal triglycerides

Feces were collected from the cages of mice that had been fed HFD for 8 weeks and triglycerides were quantified by the VUMC Lipid Core.

### Feeding behavior studies

For weekly food intake studies, individually housed animals were given *ad libitum* access to a pre- weighed amount of food which was weighed again at the end of each week. For fasting-refeeding studies, individually housed chow-fed mice were fasted for 12 h before being given *ad libitum* access to preweighed amount of HFD which was weighed hourly. For food-preference studies, individually housed chow-fed mice were given *ad libitum* access to pre-weighed amounts of both chow and HFD. At the end of seven days, chow and HFD were weighed again. Studies were conducted at room temperature unless otherwise stated.

### Imaging of GRAB_NE_ fluorescence transients in inguinal WAT

Young adult SNAP25^Δ3/Δ3^ mice or SNAP25^+/+^ littermate controls hemizygous for *Rosa26-LSL- jRGECO1A-2A-GRAB-NE* (a gift from the laboratory of Yulong Li, Peking University) and *TH*-Cre (B6.Cg-7630403G23RikTg(Th-cre)1Tmd/J, JAX) were anaesthetized with isoflurane and sacrificed via cervical dislocation. GRAB_NE_ was expressed selectively on catecholaminergic neurons with *TH*-Cre. Incisions were made dorsal-ventral at the axilla and rostral-caudal down the length of the spine. Fascia were transected and iWAT was immediately cut from connecting tissue and placed in a solution of 93 mM *N*-methyl-D-glucamine, 2.5 mM KCl, 1.2 mM NaH_2_PO_4_, 20 mM HEPES, 10 mM MgSO_4_, 0.5 mM CaCl_2_, 25 mM D-glucose, 5 mM ascorbate, 3 mM pyruvate, bubbled with 95% O_2_-5% CO_2_, pH 7.4. Inguinal WAT was mounted with small pins through the lateral edges to a silicone elastomer stage and imaged via lattice light sheet microscopy. The solution was exchanged for 124 mM NaCl, 26 mM NaHCO_3_, 1.25 mM NaH_2_PO_4_, 3 mM KCl, 2 mM CaCl_2_, 1 mM MgCl_2_, 10 mM D-glucose, bubbled with 95% O_2_-5% CO_2_. A twisted-pair stimulating electrode was placed on adjacent ganglia from T11-L1 and GRAB-NE fluorescence transients (Ex. 485nm / Em. 535nm) were recorded using stimulus trains of 100 stimuli, 100-200 mA at 20 Hz and 1ms duration. Each image was captured with 50 ms exposure time. Transient data were analyzed as the change in fluorescence intensity relative to the resting fluorescence intensity using ImageJ.

### Statistics

Data are mean ± SEM, using GraphPad Prism version 9.1.0(221) for Windows 64-bit (GraphPad Software). Analysis comparing genotype only were performed with an unpaired t-test. Analysis comparing two independent variables were performed with a two-way ANOVA or a mixed-effects model if data were missing. Multiple comparison tests were performed with the Bonferroni correction for SNAP25 genotype only and are indicated on figures by asterisks corresponding to *, P < .05; **, P < .01; ***, P < .001. Whole body energy expenditure data were analyzed using analysis of covariance (79, 80).

## Author contributions

Z.Z. conceived the initial idea for the SNAP25^Δ3/Δ3^ mouse and insulin secretion studies, designed and led the early research and GRAB_NE_ studies, conducted experiments, analyzed data, and contributed to writing the manuscript. R.P.C. designed and led subsequent research studies centered on adipose tissue browning, conducted experiments, analyzed and organized data, wrote manuscript, and made figures. R.P.C. is listed first because he initiated the investigation into the adipocyte browning, led these subsequent studies, and oversaw the final version of the manuscript and figures. A.T.G. managed the mouse colony, conducted experiments, acquired, analyzed, and managed the data. F.A. helped maintain mouse colony and helped conduct mouse studies. A.M-B. conducted the adipocyte size analysis. F.S. aided in the collection of mouse tissues. D.L. aided in the collection of mouse tissues and performed Western Blots. J.M. provided helpful insight into adipocyte innervation. J.F and Y.L. provided the GRAB_NE_ mice. S.A. designed and oversaw the studies to measure norepinephrine release using GRAB_NE_ mice. J.E.A. designed food intake and metabolic cage studies, analyzed data, and contributed to writing the manuscript. O.P.M. designed experiments, analyzed data, and wrote the manuscript. SC designed experiments and wrote the manuscript. H.H. designed experiments, wrote and organized the manuscript, performed and was responsible for the study overall.

## Acknowledgements

We would like to thank the following individuals for technical contributions to these studies: Brigitte Jia for helping with mouse tissue collection and helping conduct glucose tolerance tests in a 29.5 °C room. Justice Simonetti for helping maintain the mouse colony and conducting mouse studies, Wei Zhang and Mark Crowder for helping collect mouse tissues, Lin Zhong for conducting echocardiograms experiments and analysis, and Ginger Milne, PhD and Benlian Gao, PhD for the analysis of plasma catecholamine. Al Powers and Greg Poffenberger for conducting islet perifusion studies in the Islet Procurement & Analysis Core of the DRTC. Graphical abstract and Figure 12 were created with BioRender.com. This work was supported by NIH grants DK109204, EY10291, and NS111749 to H.H. R.P.C. was supported by National Institute of Diabetes and Digestive and Kidney Diseases grant F32 DK116520. F.S. was supported by American Diabetes Association grant 1-18-PDF-110. The Vanderbilt Mouse Metabolic Phenotyping Center (MMPC) is supported by NIH grant DK059637. We acknowledge the Vanderbilt Translational Pathology Shared Resource supported by NCI/NIH Cancer Center Support Grant 5P30 CA68485-19, the Vanderbilt Mouse Metabolic Phenotyping Center Grant 5U24DK059637- 13, and the Shared Instrumentation Grant S10 OD023475-01A1 for the Leica Bond RX. Vanderbilt University Hormone Assay and Analytical Services Core and VUMC Lipid Core are supported by NIH grants DK059637 (MMPC) and DK020593 (DRTC). The Islet Procurement & Analysis Core is supported by the Vanderbilt Diabetes Research and Training Center (P60 DK020593). VUMC Cardiovascular Physiology Core is partially supported by NIH U24 DK059637. Vanderbilt University Neurochemistry Core is supported by the Vanderbilt University Brain Institute.

## References

1. Tremmel M, Gerdtham UG, Nilsson PM, and Saha S. Economic burden of obesity: A systematic literature review. Int J Environ Res Public Health. 2017;14(4).

2. American Diabetes Association. Economic costs of diabetes in the U.S. in 2017. Diabetes Care. 2018;41(5):917–28.

3. Hird TR, Zomer E, Owen A, Chen L, Ademi Z, Magliano DJ, et al. The impact of diabetes on productivity in China. Diabetologia. 2019;62(7):1195–203.

4. Lopez C, Bendix J, and Sagynbekov K. Weighing down America: 2020 update a community approach against obesity. Milken Institute. 2020.

5. Okunogbe A, Nugent R, Spencer G, Ralston J, and Wilding J. Economic impacts of overweight and obesity: current and future estimates for eight countries. BMJ Glob Health. 2021;6(10).

6. Ceddia RP, and Collins S. A compendium of G-protein–coupled receptors and cyclic nucleotide regulation of adipose tissue metabolism and energy expenditure. Clinical Science. 2020;134(5):473–512.

7. Nauck MA, and Meier JJ. Incretin hormones: Their role in health and disease. Diabetes Obes Metab. 2018;20 Suppl 1:5–21.

8. Ahrén B. Islet G protein-coupled receptors as potential targets for treatment of type 2 diabetes. Nat Rev Drug Discov. 2009;8(5):369–85.

9. Sriram K, and Insel PA. G protein-coupled receptors as targets for approved drugs: How many targets and how many drugs? Mol Pharmacol. 2018;93(4):251–8.

10. Evans RM, and Zamponi GW. Presynaptic Ca^2+^ channels – integration centers for neuronal signaling pathways. Trends Neurosci. 2006;29(11):617–24.

11. Smrcka AV. G protein βγ subunits: central mediators of G protein-coupled receptor signaling. Cell Mol Life Sci. 2008;65(14):2191–214.

12. Zurawski Z, Yim YY, Alford S, and Hamm HE. The expanding roles and mechanisms of G protein-mediated presynaptic inhibition. J Biol Chem. 2019;294(5):1661–70.

13. Zhao Y, Fang Q, Straub SG, Lindau M, and Sharp GWG. Noradrenaline inhibits exocytosis via the G protein βγ subunit and refilling of the readily releasable granule pool via the α_i1/2_ subunit. J Physiol. 2010;588(Pt 18):3485–98.

14. Wells CA, Zurawski Z, Betke KM, Yim YY, Hyde K, Rodriguez S, et al. Gβγ inhibits exocytosis via interaction with critical residues on soluble N-ethylmaleimide-sensitive factor attachment protein-25. Mol Pharmacol. 2012;82(6):1136–49.

15. Zurawski Z, Rodriguez S, Hyde K, Alford S, and Hamm HE. Gβγ binds to the extreme C terminus of SNAP25 to mediate the action of G_i/o_-coupled G protein-coupled receptors. Mol Pharmacol. 2016;89(1):75–83.

16. Zurawski Z, Thompson Gray AD, Brady LJ, Page B, Church E, Harris NA, et al. Disabling the Gβγ-SNARE interaction disrupts GPCR-mediated presynaptic inhibition, leading to physiological and behavioral phenotypes. Sci Signal. 2019;12(569).

17. Gerachshenko T, Blackmer T, Yoon E-J, Bartleson C, Hamm HE, and Alford S. Gβγ acts at the C terminus of SNAP-25 to mediate presynaptic inhibition. Nat Neurosci. 2005;8(5):597–605.

18. Irfan M, Zurawski Z, Hamm HE, Bark C, and Stanton PK. Disabling Gβγ-SNAP-25 interaction in gene-targeted mice results in enhancement of long-term potentiation at Schaffer collateral-CA1 synapses in the hippocampus. Neuroreport. 2019;30(10):695–9.

19. Riddy DM, Delerive P, Summers RJ, Sexton PM, and Langmead CJ. G protein-coupled receptors targeting insulin resistance, obesity, and type 2 diabetes mellitus. Pharmacol Rev. 2018;70(1):39–67.

20. Thurmond DC, and Gaisano HY. Recent insights into beta-cell exocytosis in type 2 diabetes. J Mol Biol. 2020;432(5):1310–25.

21. Hwang J, and Thurmond DC. Exocytosis Proteins: Typical and Atypical Mechanisms of Action in Skeletal Muscle. Front Endocrinol (Lausanne). 2022;13:915509.

22. Goodpaster BH, and Sparks LM. Metabolic flexibility in health and disease. Cell Metab. 2017;25(5):1027–36.

23. Smith RL, Soeters MR, Wüst RCI, and Houtkooper RH. Metabolic flexibility as an adaptation to energy resources and requirements in health and disease. Endocr Rev. 2018;39(4):489–517.

24. Thorens B. Neural regulation of pancreatic islet cell mass and function. Diabetes Obes Metab. 2014;16 Suppl 1:87–95.

25. Thompson Gray AD, Simonetti J, Adegboye F, Jones C, Zurawski Z, and Hamm HE. Sexual Dimorphism in Stress-induced Hyperthermia in SNAP25Δ3 mice, a mouse model with disabled Gβγ regulation of the exocytotic fusion apparatus. Eur J Neurosci. 2020.

26. Czech MP. Mechanisms of insulin resistance related to white, beige, and brown adipocytes. Mol Metab. 2020;34:27–42.

27. Jiang H, Ding X, Cao Y, Wang H, and Zeng W. Dense intra-adipose sympathetic arborizations are essential for cold-induced beiging of mouse white adipose tissue. Cell Metab. 2017;26(4):686–92.e3.

28. Guilherme A, Pedersen DJ, Henriques F, Bedard AH, Henchey E, Kelly M, et al. Neuronal modulation of brown adipose activity through perturbation of white adipocyte lipogenesis. Mol Metab. 2018;16:116–25.

29. Henriques F, Bedard AH, Guilherme A, Kelly M, Chi J, Zhang P, et al. Single-cell RNA profiling reveals adipocyte to macrophage signaling sufficient to enhance thermogenesis. Cell Rep. 2020;32(5):107998.

30. Huesing C, Qualls-Creekmore E, Lee N, François M, Torres H, Zhang R, et al. Sympathetic innervation of inguinal white adipose tissue in the mouse. J Comp Neurol. 2021;529(7):1465–85.

31. Huesing C, Zhang R, Gummadi S, Lee N, Qualls-Creekmore E, Yu S, et al. Organization of sympathetic innervation of interscapular brown adipose tissue in the mouse. J Comp Neurol. 2021.

32. Münzberg H, Floyd E, and Chang JS. Sympathetic innervation of white adipose tissue: to beige or not to beige? Physiology (Bethesda). 2021;36(4):246–55.

33. Collins S. A heart-adipose tissue connection in the regulation of energy metabolism. Nat Rev Endocrinol. 2014;10(3):157–63.

34. Collins S. β-Adrenergic Receptors and Adipose Tissue Metabolism: Evolution of an Old Story. Annu Rev Physiol. 2022;84:1–16.

35. Cao Y, Wang H, Wang Q, Han X, and Zeng W. Three-dimensional volume fluorescence- imaging of vascular plasticity in adipose tissues. Mol Metab. 2018;14:71–81.

36. P. Wang, K.H. Loh, M. Wu, D.A. Morgan, M. Schneeberger, X. et al. A leptin–BDNF pathway regulating sympathetic innervation of adipose tissue. Nature. 2020;583(7818):839-44.

37. Kuperman Y, Weiss M, Dine J, Staikin K, Golani O, Ramot A, et al. CRFR1 in AgRP neurons modulates sympathetic nervous system activity to adapt to cold stress and fasting. Cell Metab. 2016;23(6):1185–99.

38. Madden CJ, and Morrison SF. Central nervous system circuits that control body temperature. Neurosci Lett. 2019;696:225–32.

39. Tan CL, and Knight ZA. Regulation of Body Temperature by the Nervous System. Neuron. 2018;98(1):31–48.

40. Feng J, Zhang C, Lischinsky JE, Jing M, Zhou J, Wang H, et al. A Genetically Encoded Fluorescent Sensor for Rapid and Specific In Vivo Detection of Norepinephrine. Neuron. 2019;102(4):745–61.e8.

41. Chen B-C, Legant WR, Wang K, Shao L, Milkie DE, Davidson MW, et al. Lattice light-sheet microscopy: imaging molecules to embryos at high spatiotemporal resolution. Science. 2014;346(6208):1257998.

42. Church E, Hamid E, Zurawski Z, Potcoava M, Flores-Barrera E, Caballero A, et al. Synaptic Integration of Subquantal Neurotransmission by Colocalized G Protein-Coupled Receptors in Presynaptic Terminals. J Neurosci. 2022;42(6):980–1000.

43. Ramachandran S, Rodgriguez S, Potcoava M, and Alford S. Single Calcium Channel Nanodomains Drive Presynaptic Calcium Entry at Lamprey Reticulospinal Presynaptic Terminals. J Neurosci. 2022;42(12):2385–403.

44. Barella LF, Jain S, Kimura T, and Pydi SP. Metabolic roles of G protein-coupled receptor signaling in obesity and type 2 diabetes. FEBS J. 2021;288(8):2622–44.

45. Oyler GA, Higgins GA, Hart RA, Battenberg E, Billingsley M, Bloom FE, et al. The identification of a novel synaptosomal-associated protein, SNAP-25, differentially expressed by neuronal subpopulations. J Cell Biol. 1989;109(6 Pt 1):3039-52.

46. Bark IC, Hahn KM, Ryabinin AE, and Wilson MC. Differential expression of SNAP-25 protein isoforms during divergent vesicle fusion events of neural development. Proc Natl Acad Sci U S A. 1995;92(5):1510–4.

47. Jacobsson G, Bean AJ, Scheller RH, Juntti-Berggren L, Deeney JT, Berggren PO, et al. Identification of synaptic proteins and their isoform mRNAs in compartments of pancreatic endocrine cells. Proc Natl Acad Sci U S A. 1994;91(26):12487–91.

48. Sadoul K, Lang J, Montecucco C, Weller U, Regazzi R, Catsicas S, et al. SNAP-25 is expressed in islets of Langerhans and is involved in insulin release. J Cell Biol. 1995;128(6):1019–28.

49. Jewell JL, Oh E, and Thurmond DC. Exocytosis mechanisms underlying insulin release and glucose uptake: conserved roles for Munc18c and syntaxin 4. Am J Physiol Regul Integr Comp Physiol. 2010;298(3):R517–31.

50. Zhu D, Zhang Y, Lam PPL, Dolai S, Liu Y, Cai EP, et al. Dual role of VAMP8 in regulating insulin exocytosis and islet β cell growth. Cell Metab. 2012;16(2):238–49.

51. Lam PPL, Ohno M, Dolai S, He Y, Qin T, Liang T, et al. Munc18b is a major mediator of insulin exocytosis in rat pancreatic β-cells. Diabetes. 2013;62(7):2416–28.

52. Liang T, Qin T, Kang F, Kang Y, Xie L, Zhu D, et al. SNAP23 depletion enables more SNAP25/calcium channel excitosome formation to increase insulin exocytosis in type 2 diabetes. JCI Insight. 2020;5(3).

53. Kajimura S, Spiegelman BM, and Seale P. Brown and beige fat: Physiological roles beyond heat generation. Cell Metab. 2015;22(4):546–59.

54. Tsagkaraki E, Nicoloro SM, DeSouza T, Solivan-Rivera J, Desai A, Lifshitz LM, et al. CRISPR- enhanced human adipocyte browning as cell therapy for metabolic disease. Nat Commun. 2021;12(1):6931.

55. Timmers KI, Clark AE, Omatsu-Kanbe M, Whiteheart SW, Bennett MK, Holman GD, et al. Identification of SNAP receptors in rat adipose cell membrane fractions and in SNARE complexes co-immunoprecipitated with epitope-tagged N-ethylmaleimide-sensitive fusion protein. Biochem J. 1996;320 (Pt 2)(Pt 2):429-36.

56. Araki S, Tamori Y, Kawanishi M, Shinoda H, Masugi J, Mori H, et al. Inhibition of the binding of SNAP-23 to syntaxin 4 by Munc18c. Biochem Biophys Res Commun. 1997;234(1):257–62.

57. Wang G, Witkin JW, Hao G, Bankaitis VA, Scherer PE, and Baldini G. Syndet is a novel SNAP- 25 related protein expressed in many tissues. J Cell Sci. 1997;110 (Pt 4):505–13.

58. Wong PP, Daneman N, Volchuk A, Lassam N, Wilson MC, Klip A, et al. Tissue distribution of SNAP-23 and its subcellular localization in 3T3-L1 cells. Biochem Biophys Res Commun. 1997;230(1):64–8.

59. Fischer AW, Cannon B, and Nedergaard J. Optimal housing temperatures for mice to mimic the thermal environment of humans: An experimental study. Mol Metab. 2018;7:161–70.

60. Keijer J, Li M, and Speakman JR. What is the best housing temperature to translate mouse experiments to humans? Mol Metab. 2019;25:168–76.

61. Young P, Arch JR, and Ashwell M. Brown adipose tissue in the parametrial fat pad of the mouse. FEBS Lett. 1984;167(1):10–4.

62. Burton AC, and Edholm OG. Man in a cold environment : physiological and pathological effects of exposure to low temperatures. London: Arnold; 1955.

63. Weiss B. Effects of brief exposures to cold on performance and food intake. Science. 1958;127(3296):467-8.

64. Bray GA. Reciprocal relation of food intake and sympathetic activity: experimental observations and clinical implications. Int J Obes Relat Metab Disord. 2000;24 Suppl 2:S8–17.

65. Mahú I, Barateiro A, Rial-Pensado E, Martinéz-Sánchez N, Vaz SH, Cal PMSD, et al. Brain- Sparing Sympathofacilitators Mitigate Obesity without Adverse Cardiovascular Effects. Cell Metab. 2020;31(6):1120–35.e7.

66. Dai C, Brissova M, Hang Y, Thompson C, Poffenberger G, Shostak A, et al. Islet-enriched gene expression and glucose-induced insulin secretion in human and mouse islets. Diabetologia. 2012;55(3):707–18.

67. Brissova M, Shiota M, Nicholson WE, Gannon M, Knobel SM, Piston DW, et al. Reduction in pancreatic transcription factor PDX-1 impairs glucose-stimulated insulin secretion. J Biol Chem. 2002;277(13):11225–32.

68. Rottman JN, Ni G, and Brown M. Echocardiographic evaluation of ventricular function in mice. Echocardiography. 2007;24(1):83–9.

69. Gao S, Ho D, Vatner DE, and Vatner SF. Echocardiography in Mice. Curr Protoc Mouse Biol. 2011;1:71–83.

70. Ayala JE, Bracy DP, McGuinness OP, and Wasserman DH. Considerations in the design of hyperinsulinemic-euglycemic clamps in the conscious mouse. Diabetes. 2006;55(2):390–7.

71. Ayala JE, Bracy DP, Malabanan C, James FD, Ansari T, Fueger PT, et al. Hyperinsulinemic- euglycemic clamps in conscious, unrestrained mice. J Vis Exp. 2011(57).

72. Hughey CC, Wasserman DH, Lee-Young RS, and Lantier L. Approach to assessing determinants of glucose homeostasis in the conscious mouse. Mamm Genome. 2014;25(9- 10):522–38.

73. Chi J, Wu Z, Choi CHJ, Nguyen L, Tegegne S, Ackerman SE, et al. Three-Dimensional Adipose Tissue Imaging Reveals Regional Variation in Beige Fat Biogenesis and PRDM16-Dependent Sympathetic Neurite Density. Cell Metab. 2018;27(1):226–36.e3.

74. Chi J, Crane A, Wu Z, and Cohen P. Adipo-Clear: A Tissue Clearing Method for Three- Dimensional Imaging of Adipose Tissue. J Vis Exp. 2018(137).

75. Dean KM, Roudot P, Welf ES, Danuser G, and Fiolka R. Deconvolution-free Subcellular Imaging with Axially Swept Light Sheet Microscopy. Biophys J. 2015;108(12):2807–15.

76. Hedde PN, and Gratton E. Selective plane illumination microscopy with a light sheet of uniform thickness formed by an electrically tunable lens. Microsc Res Tech. 2018;81(9):924–8.

77. Bria A, Bernaschi M, Guarrasi M, and Iannello G. Exploiting Multi-Level Parallelism for Stitching Very Large Microscopy Images. Front Neuroinform. 2019;13:41.

78. Wong JM, Malec PA, Mabrouk OS, Ro J, Dus M, and Kennedy RT. Benzoyl chloride derivatization with liquid chromatography-mass spectrometry for targeted metabolomics of neurochemicals in biological samples. J Chromatogr A. 2016;1446:78–90.

79. Kaiyala KJ, and Schwartz MW. Toward a more complete (and less controversial) understanding of energy expenditure and its role in obesity pathogenesis. Diabetes. 2011;60(1):17–23.

80. Corrigan JK, Ramachandran D, He Y, Palmer CJ, Jurczak MJ, Chen R, et al. A big-data approach to understanding metabolic rate and response to obesity in laboratory mice. Elife. 2020;9.

