## Supplemental for "Removing the Gβγ-SNAP25 brake on exocytosis enhances insulin action, promotes adipocyte browning, and protects against diet-induced obesity"

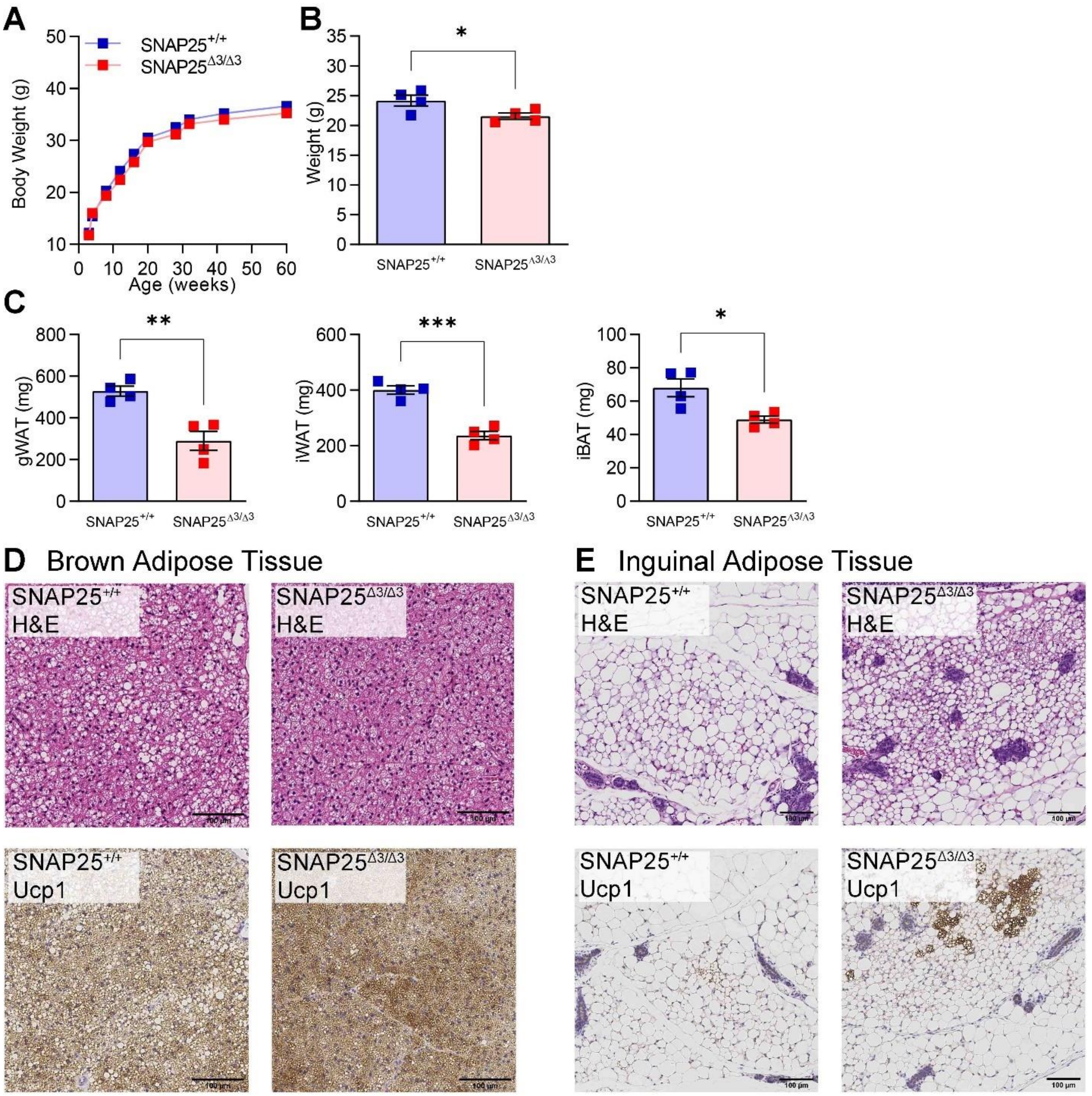
A
SNAP25+/+
SNAP25Δ3/Δ3
Body Weight (g)
Age (weeks)
B
Weight (g)
*
SNAP25+/+
SNAP25Δ3/Δ3
C
gWAT (mg)
**
iWAT (mg)
***
iBAT (mg)
*
D
Brown Adipose Tissue
SNAP25+/+
H&E
SNAP25Δ3/Δ3
H&E
SNAP25+/+
Ucp1
SNAP25Δ3/Δ3
Ucp1
E
Inguinal Adipose Tissue
SNAP25+/+
H&E
SNAP25Δ3/Δ3
H&E
SNAP25+/+
Ucp1
SNAP25Δ3/Δ3
Ucp1

**Figure S1. Body and adipose tissue weights are modestly reduced while white adipose tissue beiging is increased in chow fed, female SNAP25$^{\Delta 3/\Delta 3}$ mice**

**A.** Body weights of SNAP25$^{+/+}$ and SNAP25$^{\Delta 3/\Delta 3}$ female mice were fed standard chow diets. *N* = 12 SNAP25$^{+/+}$, 11 SNAP25$^{\Delta 3/\Delta 3}$. Analysis was performed by two-way ANOVA with repeated measures and post hoc analyses were performed using Bonferroni multiple comparisons test for SNAP25 genotype only. **B.** Body weight of a separate cohort of chow-fed mice that were euthanized at 15-weeks of age. **C.** Adipose tissue weights of female mice at 15-weeks of age. For B and C, *N* = 4 SNAP25$^{+/+}$, 4 SNAP25$^{\Delta 3/\Delta 3}$. **D.** Representative H&E- and UCP1-stained sections of iBAT. **E.** Representative H&E- and UCP1-stained sections of iWAT. Images are a representative sample from three mice from each group. * $p<0.05$, ** $p<0.01$, *** $p<0.001$. Values are expressed as mean ± SEM. All other analyses were performed using t-test.

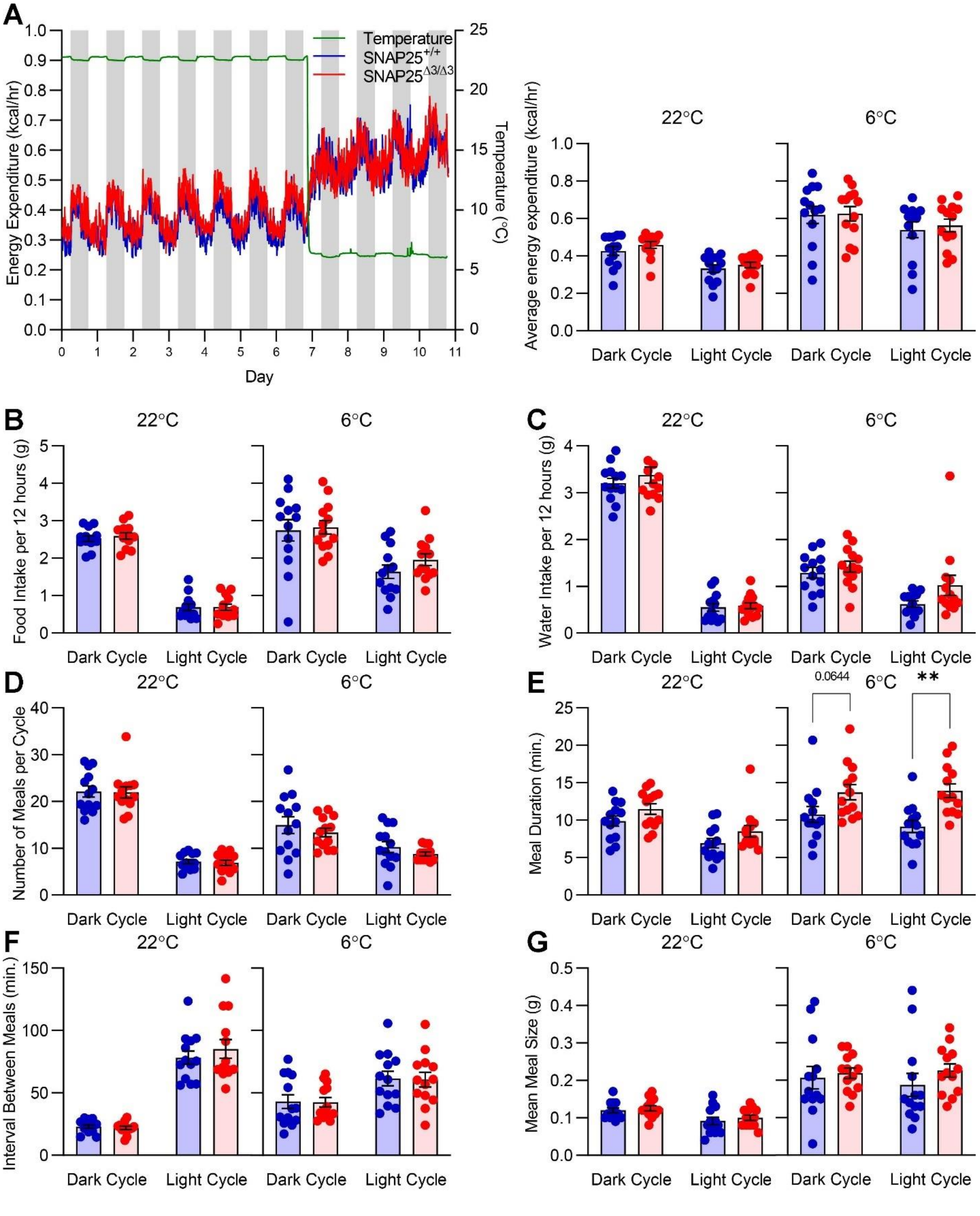
A
Temperature
SNAP25+/+
SNAP25Δ3/Δ3
Energy Expenditure (kcal/hr)
Temperature (°C)
Day
22°C
6°C
Average energy expenditure (kcal/hr)
Dark Cycle
Light Cycle
B
Food Intake per 12 hours (g)
C
Water Intake per 12 hours (g)
D
Number of Meals per Cycle
E
Meal Duration (min.)
0.0644
**
F
Interval Between Meals (min.)
G
Mean Meal Size (g)

**Figure S2. Energy expenditure at room temperature and in response to cold is unchanged in chow fed, male SNAP25$^{\Delta3/\Delta3}$ mice**

Energy balance was measured using the Promethion System at 22°C or 6°C (as indicated by green line) in 9-week-old male SNAP25$^{+/+}$ or SNAP25$^{\Delta3/\Delta3}$ mice. **A.** Energy expenditure (P = 0.1048 for the effect of genotype × light cycle interaction at 22°C). **B.** Food consumption. **C.** Water consumption (P = 0.1036 for the effect of genotype at 6°C). **D.** Number of meals per light/dark cycle. **E.** Meal duration (P = 0.0631 for the effect of genotype at 22°C, P = 0.0050 for the effect of genotype at 6°C, P = 0.0821 for the effect of genotype × light cycle interaction at 6°C). **F.** Time between the meals. **G.** Meal size (P = 0.0908 for the effect of genotype × light cycle interaction at 6°C). For B-G, *N* = 13 SNAP25$^{+/+}$, 13 SNAP25$^{\Delta3/\Delta3}$. For all figures, *N* = 13 SNAP25$^{+/+}$, 13 SNAP25$^{\Delta3/\Delta3}$. ** $p<0.01$. Values are expressed as mean ± SEM. Temperatures were analyzed separately. Analyses were performed using two-way ANOVA with each temperature analyzed separately, and post hoc analyses were performed using Bonferroni multiple comparisons test for SNAP25 genotype only and are indicated on figures.

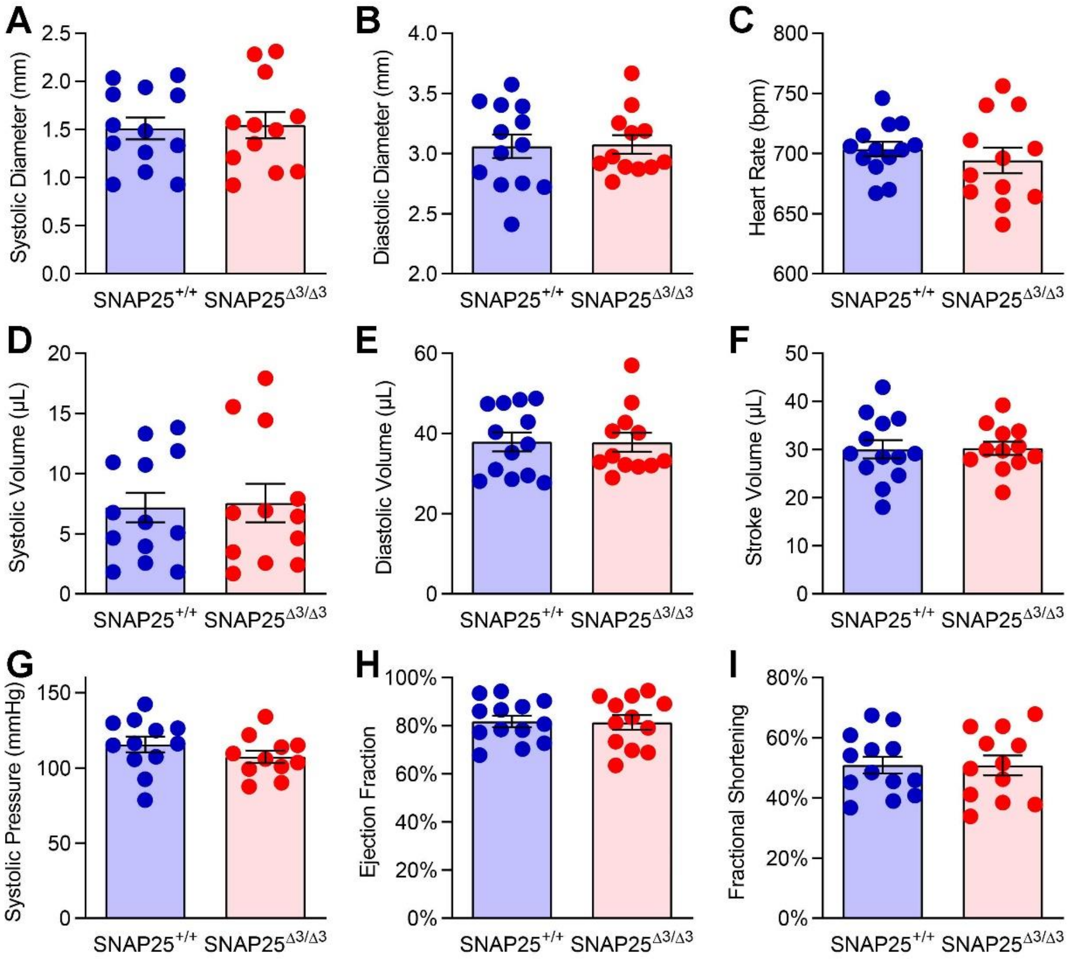


**Figure S3. Cardiovascular function is normal in male $SNAP25^{\Delta3/\Delta3}$ mice**

Cardiac parameters were measured by parasternal M-mode echocardiography in 14-week-old unanesthetized age-matched littermate $SNAP25^{+/+}$ and $SNAP25^{\Delta3/\Delta3}$ mice. No significant differences were detected between genotypes in the parameters of **A.** systolic diameter (P = 0.8563), **B.** diastolic diameter (P = 0.9025), **C.** heart rate (P = 0.4421), **D.** systolic volume (P = 0.8519), **E.** diastolic volume (P = 0.9700), **F.** stroke volume (P = 0.9326), **G.** systolic pressure (P = 0.2321), **H.** ejection fraction (P = 0.9014), **I.** fractional shortening (P = 0.9667). For all figures *N* = 13 $SNAP25^{+/+}$, 12 $SNAP25^{\Delta3/\Delta3}$. Values are expressed as mean ± SEM. Analyses were performed using t-test.

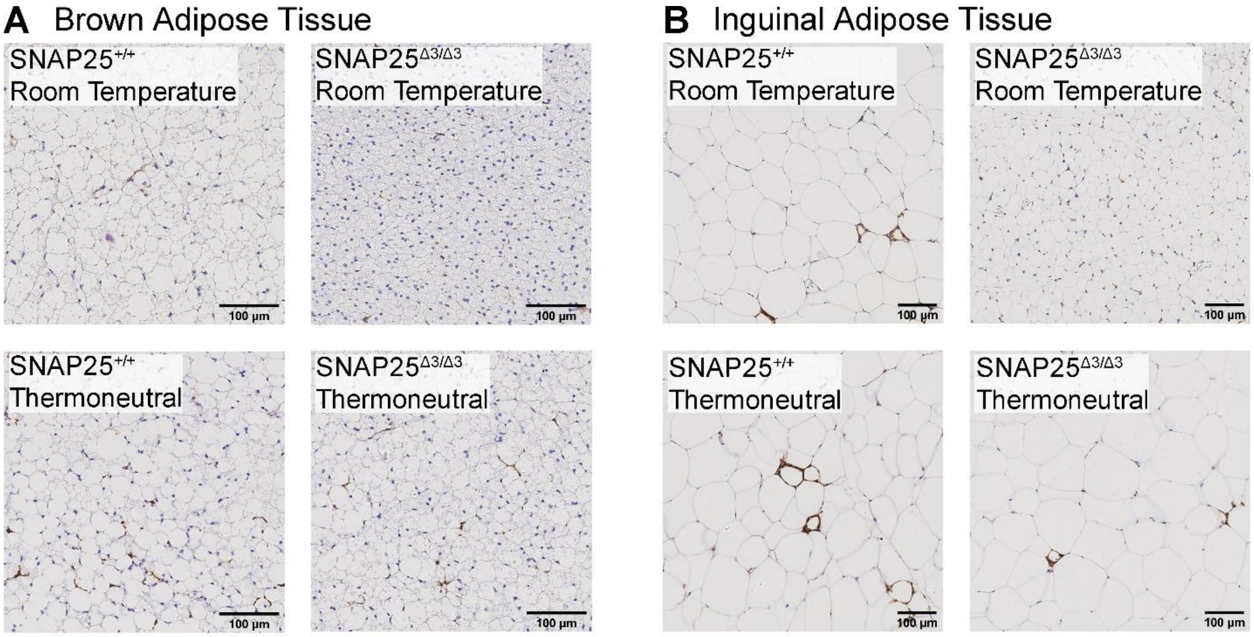


**Figure S4. Macrophage infiltration in mice with diet induced obesity**

Macrophage infiltration in HFD fed mice was assessed by F4/80 staining. **A.** Representative F4/80 staining in iBAT. **B.** Representative F4/80 staining in iWAT. Images are a representative sample from three mice from each group.

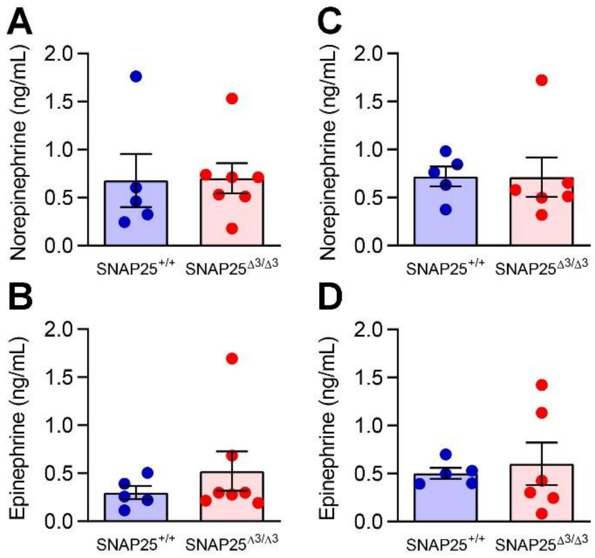


**Figure S5. Circulating catecholamine levels are unchanged in male SNAP25$^{\Delta3/\Delta3}$ mice**

Arterial plasma catecholamines obtained from chronically catheterized conscious male mice. **A.** Norepinephrine from chow-fed mice. **B.** Epinephrine from chow-fed mice. **C.** Norepinephrine from HFD-fed mice. **D.** Epinephrine from HFD-fed mice. For A-B, $N$ = 5 SNAP25$^{+/+}$, 7 SNAP25$^{\Delta3/\Delta3}$. For C-D, $N$ = 5 SNAP25$^{+/+}$, 6 SNAP25$^{\Delta3/\Delta3}$. Values are expressed as mean ± SEM. Analyses were performed using t-test.

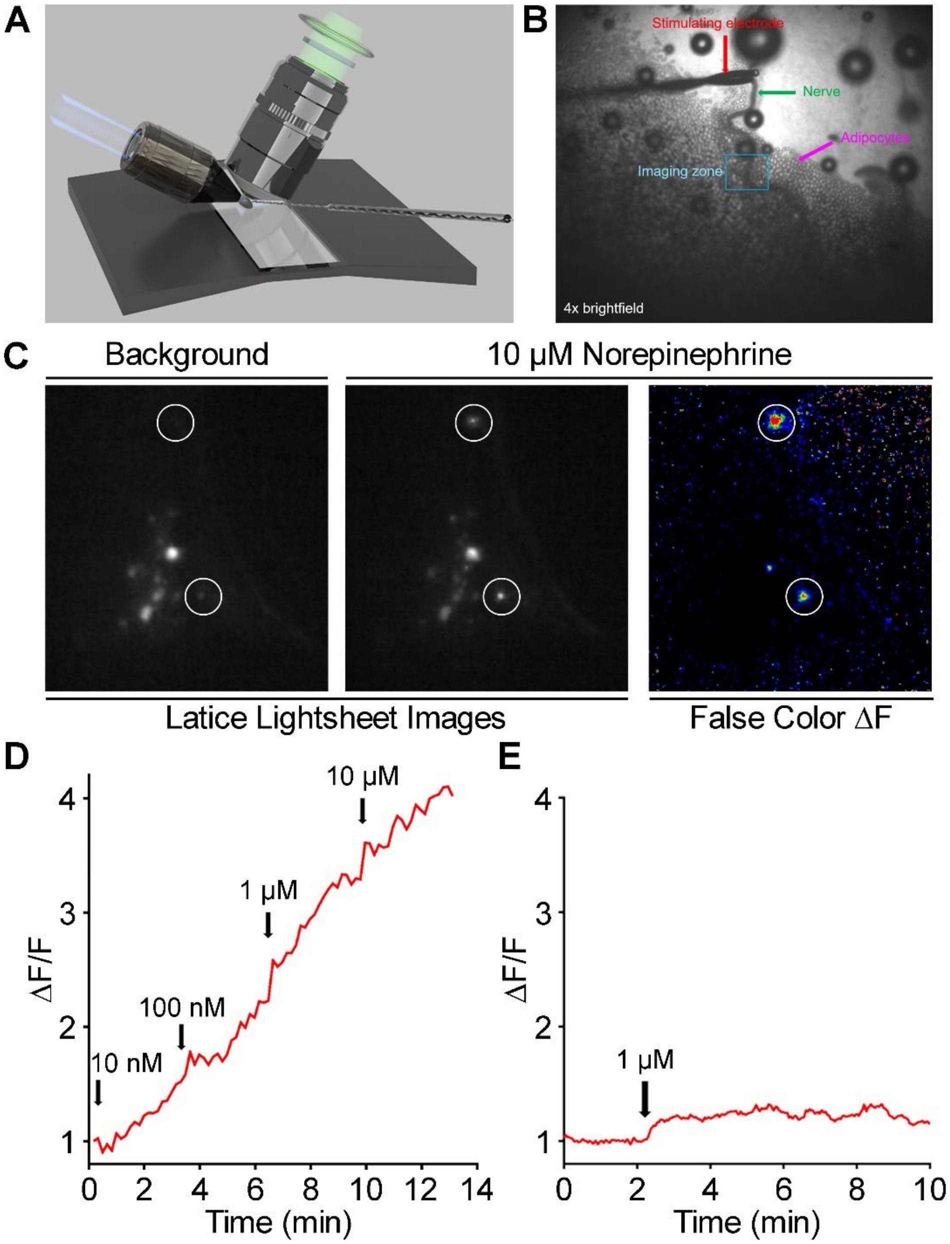
A
B
Stimulating electrode
Nerve
Adipocytes
Imaging zone
4x brightfield
C
Background
10 μM Norepinephrine
Latice Lightsheet Images
False Color ΔF
D
ΔF/F
10 nM
100 nM
1 μM
10 μM
Time (min)
E
ΔF/F
1 μM
Time (min)

**Figure S6. Fluorescence response to norepinephrine produced by $GRAB_{NE}$ expressing sympathetic neurons in inguinal white adipose tissue**

**A.** Diagram of lattice light sheet assay principle, where a pinned iWAT fat pad is stimulated by a twisted-pair electrode. Incident light sheets are shown in blue, while 535 nm fluorescence is shown in green. **B.** 4x brightfield image showing small round adipocytes (magenta arrow), T12-L1 sympathetic ganglia (green), stimulating electrode (red) and the imaging zone outlined in blue. **C.** Representative lattice light sheet microscopy images of 535 nm $GRAB_{NE}$ fluorescence from recently excised iWAT fat pads where $GRAB_{NE}$ expression is under the control of *TH*-Cre. Exogeneous norepinephrine applied to the tissue produced a large increase in fluorescence at puncta (circled), which surround the adipocytes and correspond to the sympathetic neurons. Areas of low fluorescence are depicted in blue, while areas of high fluorescence are depicted in red in the false-color heatmap. **D.** Representative dose-dependent response to exogeneous norepinephrine from a single punctum in a $GRAB_{NE}$ expressing iWAT fat pad. **E.** Representative response to exogeneous epinephrine from a single punctum in a $GRAB_{NE}$ expressing iWAT fat pad. Data are shown as values of $\Delta F/F$ where F was calculated as the mean fluorescence prior to stimulation and $\Delta F$ was fluorescence throughout the recording. In images this is shown as LUTs coding the fluorescence changes.
